# The effect of organic amendment composition on zinc and cadmium availability and uptake in wheat crops

**DOI:** 10.64898/2026.04.21.718524

**Authors:** Jill Bachelder, Julie Tolu, Lenny H.E. Winkel, Matthias Wiggenhauser, Emmanuel Frossard

**Affiliations:** ETH Zürich, Swiss Federal Institute of Technology, Department of Environment Systems Sciences (D-USYS), Institute of Agricultural Sciences (IAS), Group of Plant Nutrition, Lindau, Switzerland; ETH Zürich, Swiss Federal Institute of Technology, Department of Environment Systems Sciences (D-USYS), Institute of Biogeochemistry and Pollutant Dynamics (IBP), Group of Inorganic Environmental Geochemistry, Zürich, Switzerland; Eawag, Swiss Federal Institute of Aquatic Science and Technology, Department of Water Resources and Drinking Water (W+T), Dübendorf, Switzerland

**Author notes:** Institute of Earth Surface Dynamics, University of Lausanne, Lausanne, Switzerland (Jill Bachelder). Federal Institute of Meteorology METAS, Bern-Wabern, Switzerland (Julie Tolu). **Corresponding author:** Emmanuel Frossard, (EF).

**Keywords:** metal speciation, biogeochemistry, agriculture, cereal crops, organic fertilizers, manure, compost

## Abstract

Organic amendments provide crops with nutrients, but can also add pollutants. Yet the fate of micronutrients such as zinc (Zn) and pollutants such as cadmium (Cd) in soil-crop systems is difficult to predict because of the complexity of amendments added to soils. We performed pot and incubation experiments to determine whether the soil availability, uptake and transfer to grain of Zn and Cd in wheat (*Triticum aestivum*) are linked to the composition of amendments. Three amendments with highly diverse chemical properties, including varied organic matter (OM) degradability, were applied to a non-contaminated, arable soil. Stable isotopes of ^70^Zn and ^106^Cd were used to trace metals taken up from inputs versus soil in wheat biomass. We found the amendment most enriched in rapidly degradable OM (poultry manure) led to the highest wheat uptake of input-derived Zn i.e., 87±14 mg Zn (kg soil)^−1^. This was 2.5 times higher than input-derived Zn uptake from the most degraded amendment (compost). We did not observe an increase in soil available Zn with amendment application. Thus, biotic processes resulting from soil-plant-microbial interactions led to the increase in wheat uptake of input-derived Zn with amendment enrichment in rapidly degradable OM. Amendments led to minimal uptake of input-derived Cd in wheat and did not increase soil available Cd. Furthermore, we found no significant increase in grain Zn and Cd concentrations with amendments compared to the control. Our results highlight how amendment OM composition affects soil availability and wheat uptake of Zn and Cd with organic amendment application.

## I. Introduction

Organic amendments play an important role in agricultural systems because they allow recycling of nutrients to crops.(1–3) In particular, they can add essential metal micronutrients such as zinc (Zn), which can often be too low in agricultural soils leading to low concentrations in food products.(4,5) Amendments may also add metal contaminants like cadmium (Cd) to the soil, which may increase toxicant supply in food products leading to chronic exposure and deleterious health effects in humans.(6–8) However, the high diversity and complexity of amendments in agriculture make it very difficult to predict the fate of the metals they add.

Amendments vary widely in their pH, elemental contents, and organic matter composition,(9–13), all of which can influence Zn and Cd availability for uptake by crops.(14,15) In our recent study, we identified trends in these parameters in a wide range of amendments, including green manures, lignified crop residues and litter, monogastric farmyard manures (e.g., poultry manure), ruminant farmyard manures (e.g., cattle manure), cattle slurry, and compost.(16) We found certain amendments to be more enriched in rapidly degradable OM (e.g., plant-based amendments and monogastric farmyard manures) compared to other amendments (e.g., ruminant farmyard manure, compost). We also found large differences in water-soluble Zn and Cd speciation, which was primarily in inorganic aqueous forms in certain amendments (e.g. Zn^2+^, Cd^2+^, mainly found in ruminant farmyard manures and cattle slurries), and mainly bound to dissolved OM in other amendments (green manures, non-ruminant farmyard manures, composts). Yet, few studies have investigated the mechanisms through which amendment OM composition and water-soluble Zn and Cd speciation affect the fate of Zn and Cd in soil-plant systems.

Due to the high complexity of amendment chemical composition and plant-soil-amendment interactions, a combination of approaches is necessary to characterize mechanisms leading to solubilization and increased availability of Zn and Cd after amendment application to soil. Diffusive gradients in thin films (DGT) can be used to quantify potentially available species by sampling Zn and Cd able to diffuse in the soil solution during a short period of time (i.e., hours or days).(17) Free ion forms of Zn and Cd, which are the primary species taken up by wheat, can be predicted using chemical equilibrium modelling (e.g., WHAM VII).(18,19) Water-soluble metal-organic complexes of varying size and chemical properties can be characterized using size exclusion chromatography coupled to ultraviolet detection and inductively coupled plasma tandem mass spectrometry (SEC-UV-ICP-MS/MS).(20) Yet, soil characterization techniques can only capture Zn and Cd solubility and availability but cannot verify plant uptake. Stable isotope tracing can be used to quantify the amount of Zn and Cd from amendments that is taken up by plants, allowing differentiation from soil Zn and Cd pools.(21–23) However, no previous study has used a combination of these approaches to investigate the effect of amendment application on Zn and Cd in a soil-plant system.

We performed pot and incubation experiments to determine how amendment chemical composition affects Zn and Cd uptake by wheat. We hypothesized that application of amendments would add available species of Zn and Cd to soil, thereby leading to increased wheat uptake. We also expected the amendments enriched in fresh, rapidly degradable OM would lead to increased wheat uptake of Zn and Cd derived from soil, compared to amendments depleted in degradable OM. We expected this to occur primarily due to release of low-molecular-weight organic compounds during OM degradation, which would solubilize Zn and Cd previously immobilized in the soil solid phases. To evaluate these hypotheses, we selected three organic amendments (poultry manure, cattle manure, and compost), which were previously shown to vary in their contents of available Zn and Cd species and in OM molecular composition.(16) We applied these organic amendments and grew wheat in a soil previously labeled with stable isotopes (^70^Zn^2+, 106^Cd^2+^), allowing us to distinguish between input- and soil-derived Zn and Cd in wheat.(21) We aimed to capture the most important processes controlling the fate and dynamics of Zn and Cd in the studied soil-plant-amendment system using complementary methods. To evaluate how the amendments affected available species of Zn and Cd in the soil, we used the diffusive gradients in thin films (DGT) technique to quantify transfer of Zn and Cd via diffusion from the soil solution and solid phases to a sink.(17) To identify specific potentially available species of Zn and Cd, we characterized soil water-extractable Zn and Cd species using chemical equilibrium modelling (WHAM VII) and SEC-UV-ICP-MS/MS.(20) Finally, we investigated sorption of Zn and Cd to the soil solid phases using the multisurface modelling functions of WHAM VII.

## II. Materials and methods

### II.1 Materials

#### II.1.1 Soil

The soil used in this experiment was a Cambisol sampled from a cropped agricultural field in the canton of Jura, Switzerland. Soil was sampled at a depth of 0-20 cm in September 2020. After sampling, the soil was air-dried and a subsample was sieved to 2 mm. Analyses of pH (water, 1:2.5), cation exchange capacity and soil texture were performed by Sol-Conseil, Switzerland following standardized soil characterization methods from Agroscope Switzerland (Method S1).(24–26) Remaining soil was passed through a 6 mm plastic sieve for analyses and experiment. Water holding capacity (WHC) was determined under ambient pressure following the procedure of Agroscope.(27) Total carbon (C), nitrogen (N), and sulfur (S) concentrations were measured in samples dried at 70◦C with an elemental analyzer (Vario PYRO cube, Elementar GmbH, Germany). Diethylenetriaminepentaacetic acid (DTPA)-extractable Zn and Cd was performed following Lindsay and Norvell.(28) Iron (oxy)hydroxides, including crystalline and amorphous, was estimated using dithionite-citrate-bicarbonate extraction.(29) Total soil Zn and Cd concentrations were determined by microwave acid digestion with HF following the procedure of Imseng et al.(30) (Method S2). Total Zn and Cd analysis was performed via ICP-MS (Method S3, Table S1). The composition of the sampled soil can be found in Table 1, with some values previously reported by Schweizer et al.(31) All concentration values in Table 1 and throughout this manuscript are provided in dry weight. Soil DTPA-extractable Zn was found to be 0.8 mg Zn (kg soil)^−1^. This is in the range of 0.1-1 mg Zn (kg soil)^−1^, which can indicate limited Zn availability in soil.(32)

**Table 1.** Selected properties of soil and organic amendments. Samples of topsoil (0-20 cm) were sieved to 2 mm prior to analysis. Amendments were manually mixed and then freeze-dried. Prior to total elemental analysis, samples were ground using a tungsten-carbide ball mill. Values below the limit of detection are noted (< LOD).

| Property | Unit | Soil | Poultry manure | Cattle manure | Compost |
| --- | --- | --- | --- | --- | --- |
| pH<sup>a</sup> |  | 5.5* | 7.16 ± 0.01 | 9.0 ± 0.2 | 7.9 ± 0.3 |
| C<sup>b</sup> | g (kg)<sup>-1</sup> | 16 ± 1 | 333 ± 3 | 299 ± 7 | 152 ± 5 |
| N<sup>b</sup> | g (kg)<sup>-1</sup> | 1.7 ± 0.1 | 39 ± 1 | 25.3 ± 0.3 | 12.6 ± 0.3 |
| N-NH<sub>4</sub><sup>c</sup> | mg (kg)<sup>-1</sup> |  | 2199 | 286 | 73 |
| N-NO<sub>3</sub><sup>d</sup> | mg (kg)<sup>-1</sup> |  | < LOD | 471 | 58 |
| S<sup>b</sup> | g (kg)<sup>-1</sup> | 0.208 ± 0.001 | 3.1 ± 0.4 | 4.9 ± 0.1 | 1.5 ± 0.1 |
| DTPA-Zn<sup>e</sup> | mg (kg)<sup>-1</sup> | 0.8 ± 0.2 |  |  |  |
| DTPA-Cd<sup>e</sup> | mg (kg)<sup>-1</sup> | 0.09 ± 0.03 |  |  |  |
| Total Zn<sup>f</sup> | mg (kg)<sup>-1</sup> | 95.2 ± 0.1 | 170 ± 8 | 107 ± 2 | 141 ± 13 |
| Total Cd<sup>f</sup> | μg (kg)<sup>-1</sup> | 893 ± 10 | 361 ± 6 | 184 ± 4 | 335 ± 17 |
| Zn:Cd<sup>g</sup> |  | 106 | 471 | 581 | 419 |
| DCB-Fe<sup>h</sup> | g (kg)<sup>-1</sup> | 1.5 ± 0.1 |  |  |  |
| Clay<sup>i</sup> | g (kg)<sup>-1</sup> | 192* |  |  |  |
| Silt<sup>i</sup> | g (kg)<sup>-1</sup> | 633* |  |  |  |
| Sand<sup>i</sup> | g (kg)<sup>-1</sup> | 175* |  |  |  |
| WHC<sup>j</sup> | g (kg)<sup>-1</sup> | 320 ± 7 |  |  |  |
| CEC<sup>k</sup> | meq (100 g)<sup>-1</sup> | 13.5* |  |  |  |

<sup>a</sup>Measured in water, 1:2.5 for soil (Agroscope, 2020) and 1:50 for amendments
<sup>b</sup>Elemental analyzer, Vario PYRO cube, Elementar GmbH, Germany
<sup>c</sup>Ion chromatography via metrohm 930 Compact IC Flex with chemical suppression
<sup>d</sup>Berhelot's reaction with Agilent Cary 60 Spectrophotometer (Searle, 1984); LOD = 1 mg (L)<sup>-1</sup>
<sup>e</sup>Lindsay and Norvell, 1978
<sup>f</sup>Acid digestion (HF for soil, HNO<sub>3</sub> for amendments); see Methods S2 and S3
<sup>g</sup>By mass
<sup>h</sup>Dithionite-citrate-bicarbonate extractable iron (Mehra and Jackson, 1960)
<sup>i</sup>Gravimetric analysis (Agroscope, 2020)
<sup>j</sup>Water-holding capacity (Agroscope, 2020)
<sup>k</sup>Cation exchange capacity (Agroscope, 2020)
\*Measurement performed by Sol Conseil, Gland, Switzerland

#### II.1.2 Organic amendments

Poultry manure was sampled from Agrovet-Strickhof (Zurich, Switzerland). Cattle manure was sampled from Agroscope-Reckenholz (Zurich, Switzerland). Compost was sampled from the Biomassehof AG composting plant (Winterthur, Switzerland). Further sample details are provided in Table S2 and Note S1. Prior to analysis and soil amendment, all amendment samples were freeze-dried and finely ground using a Quiagen TissueLyzer ball mill with tungsten-carbide beakers. Total elemental concentrations (C, N, S, Zn, Cd) were analyzed as described for soil (see II.1.1 and Methods S2-3). Amendment pH was measured in ultrapure water with a solid-liquid ratio of 1:50 (Metrohm, Biotrode). Mineral N was determined by quantifying N-NO_3_^−^ and N-NH_4_^+^ in 0.01 M CaCl_2_ extracts (solid-liquid ratio of 1:20) via ion chromatography (Metrohm 930 Compact IC Flex with chemical suppression) and Berhelot’s reaction(33) (Agilent Cary 60 Spectrophotometer), respectively. For characterization of water-soluble Zn and Cd bound to dissolved organic matter, original freeze-dried amendments were extracted with ultrapure water (>18.2 MΩ, solid-liquid ratio of 1:50 for poultry manure and 1:10 for cattle manure and compost) in pre-weighed 15-mL polypropylene falcon tubes (VWR Metal-Free Centrifuge Tubes). Details of analysis are provided in section II.5.

### II.2 Experimental design

#### II.2.1 Pot experiment

A one-factorial pot experiment was performed in which wheat (cultivar Bobwhite) was grown under six treatments. An overview of the experimental setup can be found in Figure S1. The fraction of metals in the soil available for plants was labeled with ^70^Zn (97.4 atom% abundance, CortecNet SA) and ^106^Cd isotopes (99.5 atom% abundance, CortecNet SA). To achieve this, we digested solid metal blocks of ^70^Zn and ^106^Cd in 8 M HNO_3_ (double-distilled from Supelco Emsure grade). We diluted this solution to concentrations of 40 mg (L)^−1 70^Zn and 0.4 mg (L)^−1 106^Cd. To 1 kg soil, we added 200 μg of ^70^Zn as aqueous Zn(NO_3_)_2_ and 2 μg of ^106^Cd as aqueous Cd(NO_3_)_2_. After labeling, soil was moistened to 30% WHC, mixed for 2 minutes, and further moistened to 192 g H_2_O (kg soil)^−1^, or 60% WHC. To equilibrate the added isotope with the soil matrix, pots were stored under dark conditions in a growth chamber (60% relative humidity, 18ºC) for 12 weeks before the start of the pot experiment.(34)

Six treatments were applied to the soil, including no Zn input (control), mineral Zn input as aqueous ZnSO_4_, ammonium input as aqueous (NH_4_)_2_SO_4_, and three organic amendments (poultry manure, cattle manure, and compost). These three organic amendments were selected to cover variability in OM composition based on our previous study.(16) Poultry manure represented the amendment with the most degradable OM. Cattle manure was depleted in rapidly degradable OM compared to the poultry manure. Compost represented the amendment most depleted in rapidly degradable OM. Organic amendments were applied to the soil at a constant rate of 2 g C (kg soil)^−1^ as shown in Table S3. This resulted in a Zn application rate of 0.7-1.8 mg Zn (kg soil)^−1^ and Cd application rate of 1.2-4.4 μg Cd (kg soil)^−1^. For the mineral Zn treatment, ZnSO_4_ was applied as a solution of 400 mg Zn (L)^−1^, resulting in an application rate of 1.5 mg Zn (kg soil)^−1^. For the ammonium treatment, (NH_4_)_2_SO_4_ was applied as an aqueous solution of 224 mg N-NH_4_ (L)^−1^. In addition to these inputs, a basal Zn-free nutrient solution was applied to all pots (Table S3). As N can be an important driver of Zn uptake, mineral N in the basal nutrient solution was adapted to maintain a constant application rate of 100 mg mineral N (kg soil)^−1^ in all treatments. Each treatment was replicated four times. Greenhouse growing conditions, plant protection measures, and details of harvest are outlined in Method S4. Half the pots were harvested at the end of tillering (harvest 1, eight weeks after sowing), while the remaining plants were harvested at full maturity (harvest 2, 19 weeks after sowing).

#### II.2.2 Incubation experiment

A parallel incubation experiment allowed investigation of CO_2_ emissions which we assumed were due to microbial respiration. Cups filled with 21 g soil were placed in sealed jars after treatment application. Treatments were applied as shown in Table S4. No basal nutrient solution was applied.

### II.3 Soil analyses

Soil microbial respiration was measured following the procedure of Alef et al.(35) Produced CO_2_ was trapped in 0.1 M NaOH and quantified via back-titration with 0.1 M HCl (Titrisol, Merck). Respiration was measured for nine weeks.

Soil was sampled at harvest 1 (8 weeks after sowing) by manually separating roots from soil. The pH of the moist soil was measured in ultrapure water with a solid-liquid ratio of 1:2.5 (pH meter from Thermo Scientific, Orion Star A111). Soil samples were extracted with ultrapure water (>18.2 MΩ, 1:10 weight-to-volume ratio) in pre-weighed 50-mL polypropylene falcon tubes (VWR Metal Free Centrifuge Tubes). Extracts were centrifuged (15 minutes, 3000 rpm), and filtered (BGB syringe filter, 0.45 μm pore size, Nylon 66). Following extraction, soil was dried in the original extraction tubes (70 °C, 24 h) and dry weight was measured. Aliquots of 10 mL water extracts of soils and amendments were frozen in 15-mL polypropylene tubes (Grenier) and analyzed for dissolved organic carbon (DOC) and dissolved total nitrogen (DTN) (Shimadzu, TOC-L, TNM-L). For soil, aliquots of 5 mL were frozen and measured for total anion concentrations (N-NO_3_^−^, SO_4^2-^_, F^−^, Cl^−^) via ion chromatography using a Metrohm 930 Compact Ion Chromatography Flex. In addition, soil was extracted with 0.43 M HNO_3_ (1:10 weight-to-volume ratio) in acid-cleaned 15 mL polypropylene tubes (Grenier), followed by centrifugation and filtration.(36)

Diffusive gradients in thin films (DGT® research, LSLM-NP, Chelex binding layer), conditioned in 0.01 M NaNO_3_, were used to passively sample Zn and Cd able to diffuse in the soil.(37) While classical extractants such as water determine the available Zn and Cd in soils based on a chemical equilibrium between the extractant and the soil, DGT acts as a passive sink for Zn and Cd to simulate root uptake.(17,38) Briefly, the equivalent of 50 g soil was weighed in acid-cleaned plastic beakers, saturated to 100 % WHC, covered in parafilm, and incubated at 24 °C for 24 hours. DGT samplers were rinsed with ultrapure water, and a thick paste of the saturated soil was smeared on each sampler, followed by deployment into the beakers. Beakers were again covered in parafilm and incubated at 24 °C for 72 hours, then resins were retrieved and trace elements were eluted from the resins using 1 mL of 1 M HNO_3_ (double-distilled from Supelco Emsure grade).

### II.4 Plant analyses

Prior to analysis, dried plant organs were ground using a ball mill with tungsten-carbide beakers (Quiagen, TissueLyzer). Total carbon and nitrogen were measured (Vario PYRO cube, Elementar GmbH, Germany) in addition to metals (see II.5).

### II.5 Metal analyses

For plants, total element concentrations were determined by microwave acid digestion (Method S2) and ICP-MS/MS (Method S3, Note S2). For each harvested wheat plant, the whole-plant uptake of Zn and Cd was calculated (i.e., in all above- and below-ground biomass). This allowed determination of the amount of Zn and Cd accessed by the plant per kilogram of soil. An aliquot of the plant digest was purified for measurement of Zn and Cd isotope ratios using anion exchange solid-phase extraction following Method S5.(22,23,39) The quantities of Zn and Cd derived from the input (i.e., from the amendment or the ZnSO_4_) and derived from the soil were calculated as shown in Method S5.(21,22) The quantities of Zn or Cd derived from the input are noted as input-derived Zn or Cd in the rest of the text and expressed in mg or μg (kg soil)^−1^. Similarly, the quantities of Zn or Cd derived from the soil are noted as soil-derived Zn and Cd and expressed in mg or μg (kg soil)^−1^. The input-derived data were used to calculate the fertilizer use efficiency i.e., the percentage of total applied Zn and Cd quantified in plant biomass.

For soil, concentrations of Zn and Cd in DGT extracts were analyzed using ICP-MS/MS following Method S3. Effective concentrations of Zn and Cd at the soil-sampler interface (C_DGT_) were calculated as described previously.(40) Low concentrations of DGT-extractable Zn and Cd were too similar to the contamination of Zn and Cd introduced in the isotope purification procedure, thus ^70^Zn/^66^Zn and ^106^Cd/^111^Cd could not be measured in the DGT extracts.

To investigate association of soil water-extractable Zn and Cd with dissolved organic matter, SEC-UV-ICP-MS/MS was used to detect three size and chemical fractions. As a quality check, concentrations of water-soluble Zn and Cd were quantified in soil water extracts acidified to 0.3M HNO_3_, using NIST 1643f as a quality control standard (Table S5). While low concentrations of Zn were found in the soil extracts, this was still deemed acceptable for SEC-UV-ICP-MS/MS analyses (see Note S3 and Table S6). For both soil samples and amendments, aliquots of 1.5 mL of water extracts were transferred to amber borosilicate HPLC vials, stored at 4°C, and measured within 1-2 days by SEC-UV-ICP-MS/MS as described in Method S6 and previously in Tolu et al. and Bachelder et al.(16,20) Three fractions of Zn and Cd bound to DOM were determined following the original study that optimized the SEC-UV-ICP-MS/MS method (fraction 1 = “(oxy)hydroxide nanoparticles”, fraction 2 = “higher-molecular-weight OM”, fraction 3 = “lower-molecular-weight OM”).(20) Definition of these fractions was based on the peaks of UV (a proxy for OM), the peaks of monitored element (i.e., P, S, Fe, Zn, and Cd), and the elution of DOM reference materials from the International Humic Substances Society (Note S4, Figures S2-3).

To complement SEC analysis of DOM-bound Zn and Cd, multisurface modelling (WHAM VII) was used to investigate water-soluble Zn and Cd species in free ion forms (i.e., Zn^2+^, Cd^2+^) versus bound to DOM. We also used WHAM to predict Zn and Cd likely to be sorbed with low affinity to the soil solid phases i.e., that can be desorbed rapidly.(18,41,42) To achieve this, the speciation of Zn and Cd was determined in 0.43 M HNO_3_, an extractant used for estimation of sorbed species via multisurface modelling.(36) Details of equilibrium modelling are provided in Method S7, Figures S4-5, Table S7.

### II.6 Statistics

All statistical analyses were performed using R studio (version 2022.02.3+492). Significant differences between treatments were calculated using analysis of variance (ANOVA) and Tukey’s honest significant difference post hoc tests. An evaluation of the normal distribution of residuals was performed using the Shapiro-Wilk test and the equality of variances was tested using a Levene Test. If these requirements were not fulfilled, inverse, log_10_, and square root transformed data was tested. If the transformed data still failed these tests, a non-parametric Kruskal-Wallis test was used instead of the ANOVA test to determine statistical differences.

## III. Results

### III.1 Microbial respiration

For all treatments, microbial respiration increased throughout the experiment, plateauing 8 weeks after the treatments were mixed into the soil (Figure S6). Poultry manure, cattle manure, and compost increased microbial respiration compared to the control, mineral Zn, and ammonium treatments. After 9 weeks of incubation, 750±40 mg C (kg soil)^−1^ was respired with poultry manure application, the equivalent of 50% of the applied 1.5 g C (kg soil)^−1^ (Table S4).

### III.2 Wheat growth and uptake of Zn and Cd

Poultry manure application led to the highest wheat biomass at harvest 2 (Figure S7). No other treatment showed a significant difference from the control value of 10±1 g (kg soil)^−1^. Biomass weight and concentrations of C, N, Zn and Cd in root, shoot, and grain can be found in Table S8.

Plant uptake represents the mass of Zn and Cd transferred from the soil to the wheat plant (Figure 1A,B). At harvest 1, plant Zn uptake was highest with mineral Zn and poultry manure (65-120 µg Zn (kg soil)^−1^) followed by cattle manure and compost (52-108 µg Zn (kg soil)^−1^). At harvest 2, total Zn uptake increased by a factor of two compared to harvest 1. At harvest 2, all treatments except compost and mineral Zn were significantly greater than the control value of 168±11 µg Zn (kg soil)^−1^.

**Figure 1.**
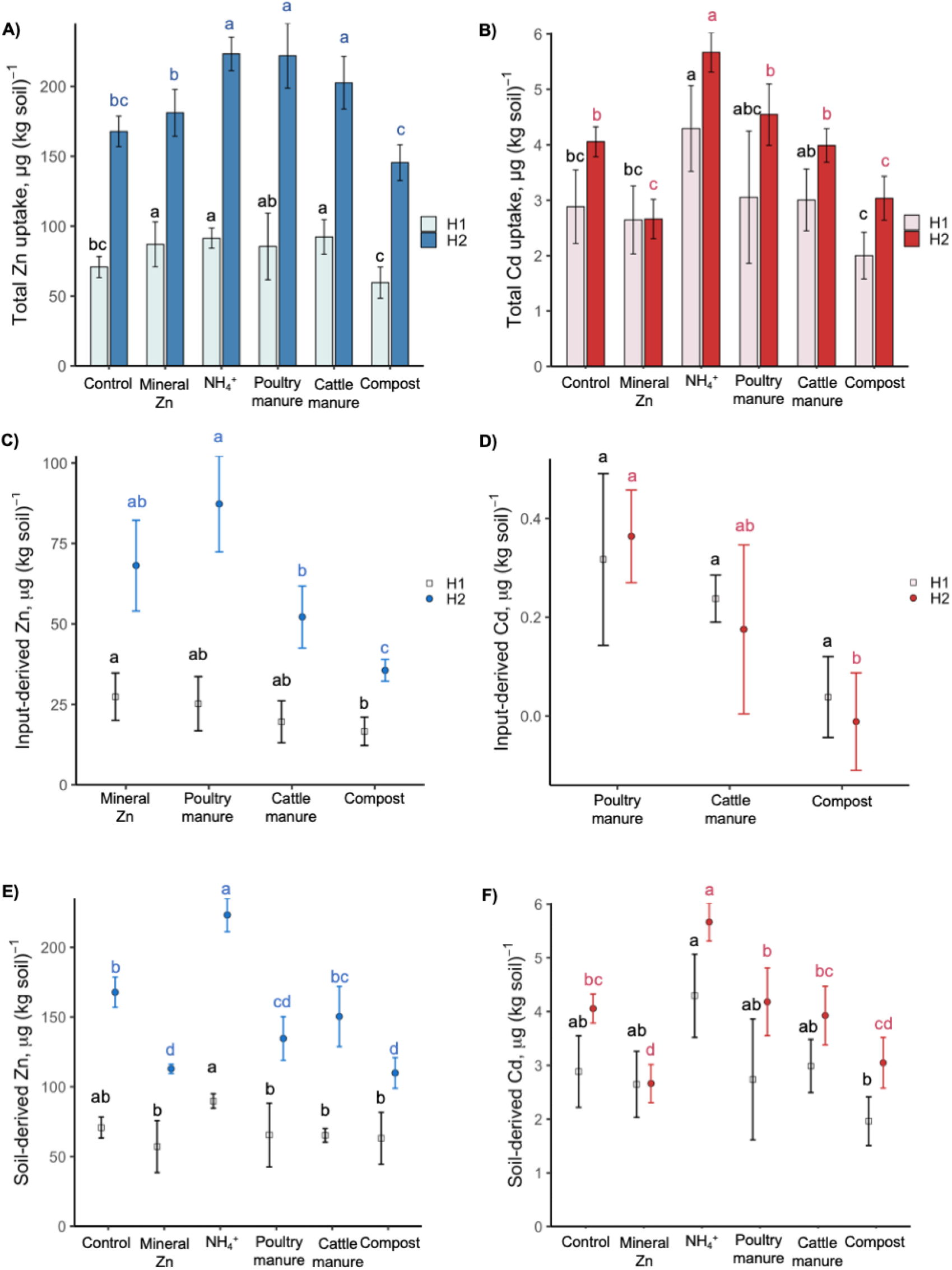
Total uptake and isotope tracing of Zn and Cd in wheat plants. Total uptake of A) Zn and B) Cd in wheat were derived from multiplying elemental concentrations in roots, stem+leaves, and grains by plant biomass. Input-derived C) Zn and D) Cd taken up by wheat was determined by isotope labeling. Soil-derived E) Zn and F) Cd taken up by wheat was also determined. Treatments under which wheat was grown included no Zn fertilizer (control), mineral Zn applied as ZnSO_4_, ammonium applied as (NH_4_)_2_SO_4_, and three organic amendments. Harvests were completed at the end of tillering (harvest 1; H1) and at full maturity (harvest 2; H2). Error bars show ±1 standard deviation of the mean, calculated from n=4 experimental (pot) replicates. Black lowercase letters a-c indicate statistical differences between treatments at harvest 1. Colorful lowercase letters a-d indicate statistical differences between treatments at harvest 2.

Plant uptake of input- and soil-derived Zn and Cd is provided in Figure 1C-F. At both harvests, soil-derived Zn was approximately two times higher than input-derived Zn. At harvest 1, the input-derived Zn from mineral Zn and poultry manure application reached 27±7 µg Zn (kg soil)^−1^ and 25±8 µg Zn (kg soil)^−1^, respectively (Fig. 1C). At harvest 2, these values increased three-fold for mineral Zn and poultry manure to 68±14 µg Zn (kg soil)^−1^ and 87±14 µg Zn (kg soil)^−1^, respectively. At harvest 2, cattle manure and compost led only to a two-fold increase in input-derived Zn compared to harvest 1 and were significantly lower than poultry manure. Ammonium application significantly increased soil-derived Zn compared to the control, to 223±12 µg Zn (kg soil)^−1^ (Fig. 1E). Organic amendments did not increase wheat uptake of soil-derived Zn at harvests 1 or 2, compared to the control. Mineral Zn, poultry manure, and compost application led to decreased soil-derived Zn compared to the control.

At harvest 1, whole-plant Cd uptake was highest with ammonium input (4.3±0.8 µg Cd (kg soil)^−1^), with organic amendments ranging 1.7-4.8 µg Cd (kg soil)^−1^ (Figure 1B). Contrary to Zn, Cd uptake showed little increase from harvest 1 to 2. Overall, input-derived Cd values were very low and showed high variability (Figure 1D). Poultry manure showed the highest input-derived Cd values, with 0.36±0.09 µg Cd (kg soil)^−1^ at harvest 2. In contrast to Zn, no increase in input-derived Cd uptake was observed at harvest 2 compared to harvest 1. For soil-derived Cd, the ammonium treatment significantly increased whole-plant uptake of soil-derived Cd at harvest 2 compared to the control (Fig. 1F). The mineral Zn treatment led to decreased soil-derived Cd at harvest 2 compared to the control.

Using values of input-derived Zn and Cd uptake in plant biomass at harvest 2 (Fig. 1C, D) and original application rates (Table S3), we further calculated the fertilizer use efficiency (Table S9). Poultry and cattle manure showed the highest use efficiency of Zn, with 8±1% and 7±1%, respectively. Thus, 7-8% of the Zn applied with these inputs reached the plant, while 92-93% of the Zn remained in the soil. With mineral Zn, use efficiency was significantly smaller (5±1%), while the compost Zn use efficiency was lowest of all inputs (2.0±0.2%). The Cd use efficiency showed high variability, likely due to low input-derived Cd values (Table S9). Similar to Zn, poultry manure application led to the highest Cd use efficiency among the organic amendments. The Cd use efficiency with cattle manure and compost was very close to zero.

### III.3 Concentrations of Zn and Cd in wheat grains

No organic amendments significantly affected grain concentrations of Zn compared to the 29±3 mg Zn (kg grain)^−1^ found in the control treatment (Figure 2A). Compared to the control, mineral Zn and ammonium application led to increased Zn by a factor of 1.3. These values were close to the threshold set by Harvest Plus as a target for Zn biofortification of wheat.(43) Similar to Zn, no organic amendments significantly affected grain concentrations of Cd compared to the control value of 0.13±0.03 mg Cd (kg grain)^−1^ (Figure 2B). Contrary to Zn, neither the mineral Zn nor ammonium treatments significantly affected Cd grain compared to the control.

**Figure 2.**
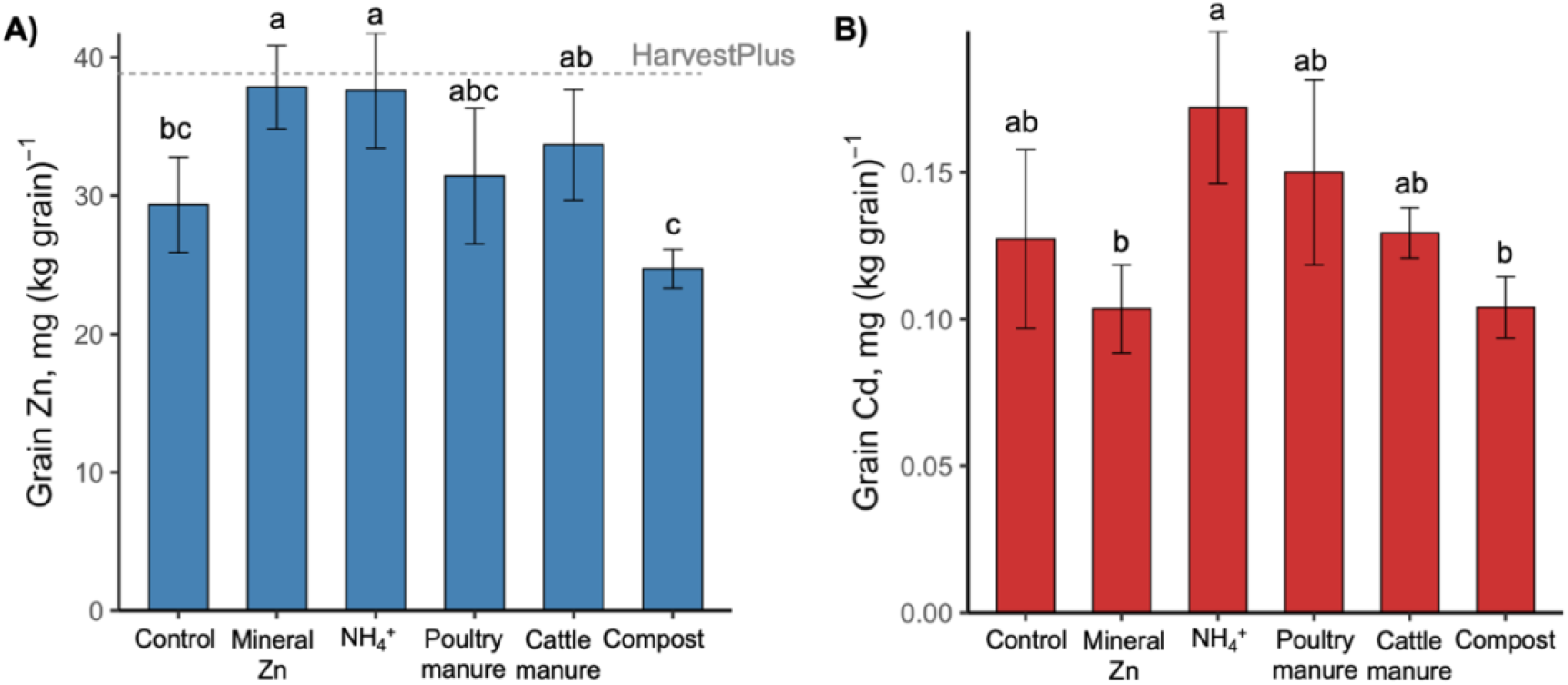
Grain concentrations of A) Zn and B) Cd in wheat-growth pot experiment. Panel A) also includes the HarvestPlus recommended target Zn concentration in wheat grains of 38 mg (kg grain)^−1^.(44) Grains were harvested at full maturity (19 weeks after sowing). Treatments under which wheat was grown included no Zn fertilizer (control), mineral Zn applied as ZnSO_4_, ammonium applied as (NH_4_)_2_SO_4_, and three organic amendments. Error bars show±1 standard deviation around the mean, calculated from n=4 experimental (pot) replicates. Lowercase letters (a-c) indicate statistical differences between treatments. Concentrations are also presented in Table S8.

### III.4 Characterization of Zn and Cd in soil pools at harvest 1

As a proxy for soil available Zn and Cd, DGT-extractable Zn and Cd were measured. Mineral Zn significantly increased the concentrations of DGT-Zn i.e., to 1.3±0.2 µg Zn L^−1^ compared to 0.5±0.2 µg Zn L^−1^ in the control (Figure 3A). Ammonium application significantly increased DGT-Zn to 1.4±0.5 µg Zn L^−1^ and increased DGT-Cd to 0.27±0.08 µg Cd L^−1^, compared to 0.09±0.04 µg Cd L^−1^ in the control (Figure 3B). This corresponded to decreased pH with ammonium application (Figure S8-A, Note S5). No organic amendments significantly changed the DGT-Zn or DGT-Cd compared to the control.

**Figure 3.**
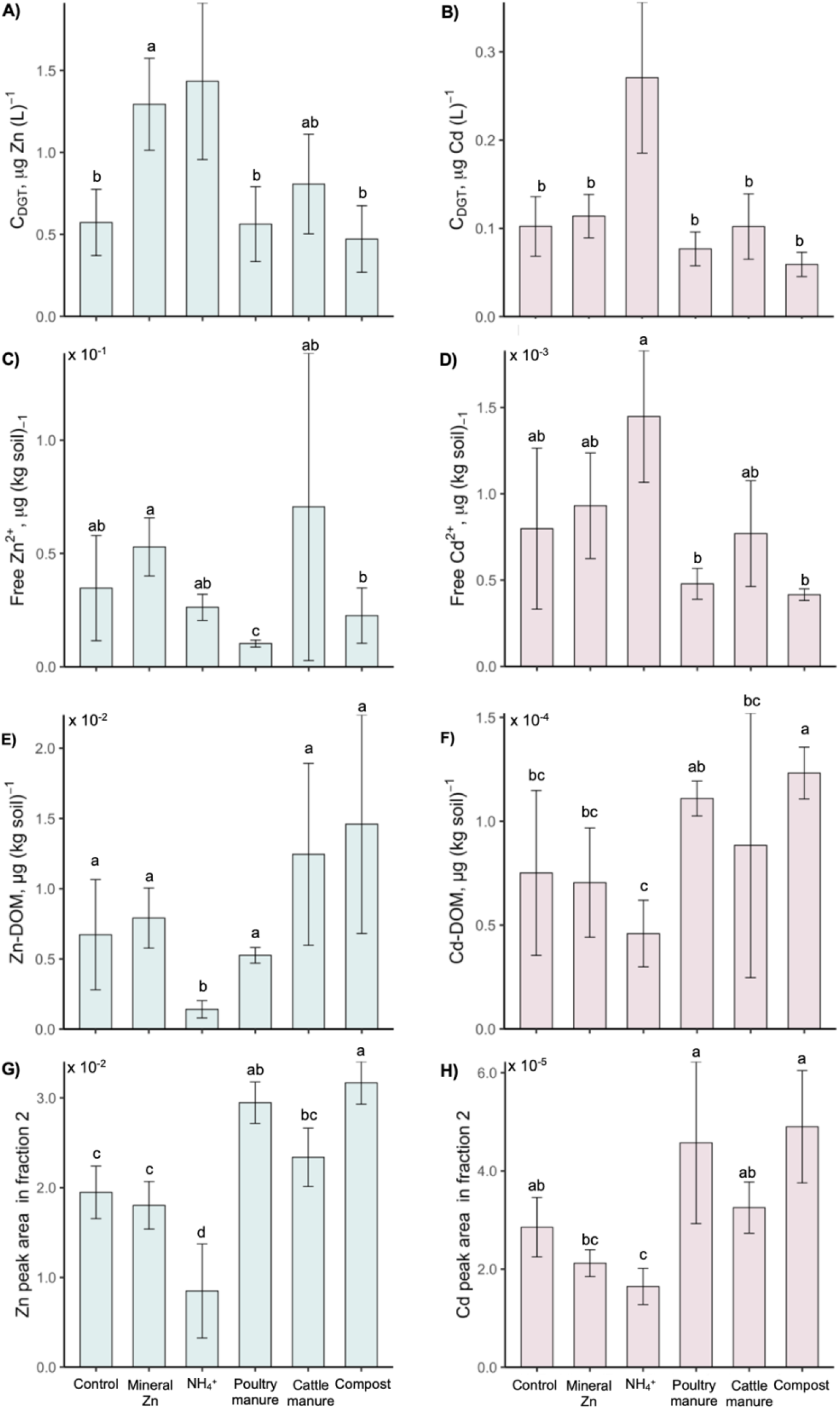
Characterization of Zn and Cd species in soil sampled from a wheat-growth pot experiment. Soil was destructively sampled and characterized at the end of tillering (harvest 1). Concentrations measured by diffusive gradients in thin film (C_DGT_) are shown for A) Zn and B) Cd. Water-extractable Zn and Cd speciation was determined using WHAM VII, including C) free Zn^2+^, D) free Cd^2+^, E) Zn bound do dissolved organic matter (DOM), and F) Cd bound to DOM. Modelling was performed with reactive DOM values of 65%. Binding of water-extractable Zn and Cd to dissolved organic matter, as measured by size exclusion chromatography coupled to ultraviolet detection and inductively coupled plasma tandem mass spectrometry (SEC-UV-ICP-MS/MS), is shown for G) Zn and H) Cd in SEC fraction F2 i.e., higher-molecular-weight dissolved organic matter. Data is presented as total peak area in each fraction as determined by peak deconvolution with the software OriginPro2021. Treatments under which wheat was grown included no Zn fertilizer (control), mineral Zn applied as aqueous ZnSO_4_, an ammonium (NH_4_^+^) treatment applied as (NH_4_)_2_SO_4_, and three organic amendments. Error bars show±1 standard deviation around the mean, calculated from n=4 experimental (pot) replicates. Lowercase letters (a-d) indicate statistical differences between treatments.

The data obtained for Zn and Cd speciation by SEC-UV-ICP-MS/MS and chemical equilibrium modelling (WHAM VII) are presented in Figure 3C-H. WHAM-predicted free Zn^2+^ ranged from 9 to 170 ng (kg soil)^−1^ and free Cd^2+^ ranged from 0.3 to 1.8 ng (kg soil)^−1^ (Figure 3C,D). Contrary to DGT-extractable pools, free Zn^2+^ did not significantly increase with mineral Zn and neither free Zn^2+^ nor free Cd^2+^ significantly increased with ammonium application, compared to the control. The WHAM-calculated water-extractable species bound to DOM ranged from 0.5 to 26 ng (kg soil)^−1^ for Zn and 0.02 to 0.17 ng (kg soil)^−1^ for Cd (Figure 3E,F). For both Zn and Cd, DOM-bound species were significantly higher with poultry manure and compost treatments and significantly lower with ammonium treatment compared to the control.

With SEC-UV-ICP-MS/MS analysis, between 6 and 71% of total water-extractable Zn and Cd could be detected (Figure S9) and was classified into three size and chemical fractions i.e., (oxy)hydroxide nanoparticles (fraction F1), higher-molecular-weight dissolved organic matter (fraction F2), and lower-molecular-weight dissolved organic matter (fraction F3). Overall, Zn and Cd were found primarily in fractions F1 and F2 (Figure S10-E,F). Very little Zn and Cd was found in fraction F3. In contrast, fraction F3 made up >70% of water-extractable Zn in the original poultry manure (Figure S11).

Otherwise, DOM-bound Zn and Cd were primarily present in fraction F2 in the original organic amendments (Figure S11). The results obtained with SEC-UV-ICP-MS/MS demonstrated a clear effect of organic amendments on water-extractable Zn and Cd bound to higher-molecular-weight DOM (F2) as compared to the control. Indeed, poultry manure and compost application significantly increased the amount of Zn in fraction F2 compared to the control (Figure 3G). Ammonium application decreased both Zn and Cd in fraction F2 (Figure 3G,H). Correlation analyses revealed that Zn and Cd in fraction F2 negatively correlated with DGT-Zn and DGT-Cd concentrations (Figure S12, Table S10).

To gain insight into Zn and Cd sorption onto the soil solid phases, the 0.43 M HNO_3_-extractable pool was used for WHAM modelling. Zn in this pool was approximately 3.8 mg (kg soil)^−1^ i.e., 2.7 times higher than the Zn application rate of approximately 1.5 mg (kg soil)^−1^. Cd was approximately 0.17 mg (kg soil)^−1^, which was >80 times higher than the Cd application rate. Overall, this pool accounted for 2.6-4.3 % of total Zn and 14-19% of the total Cd in the soil. WHAM revealed >70% of 0.43 M HNO_3_-extractable Zn and Cd sorbed to SOM and metal oxides (Figure S13). Mineral Zn, poultry manure, and compost application led to increased WHAM-calculated Zn sorption to the soil solid phases, mainly SOM and metal oxides, compared to the control (Figure S13-C). In contrast, ammonium application decreased 0.43 M HNO_3_-extractable Zn and Cd sorbed to the soil solid phases compared to the control (Figure S13-C,D). Overall, WHAM-calculated Cd sorption followed similar trends as Zn, though Cd was generally more associated with metal oxides and less associated with SOM compared to Zn (Figure S13).

## IV. Discussion

### IV.1 Amendments enriched in rapidly degradable OM increased wheat uptake of input-derived Zn but not Cd

The finding that microbial respiration followed organic amendment enrichment in rapidly degradable OM i.e., poultry manure > cattle manure > compost (Figure S6), confirmed the hypotheses of our previous characterization study.(16) As organic amendment samples may have contained carbonates, a small portion of the respired C could be due to carbonate dissolution at circumneutral pH, though it is unlikely that this contributed to significantly to values measured.(45,46) Isotope source tracing revealed that organic amendment enrichment in rapidly degradable OM leads to increased input-derived Zn (Figure 1C). Furthermore, Zn use efficiency obtained with the organic inputs more enriched in rapidly degradable OM (poultry and cattle manures) was higher than use efficiency with the highly available Zn form of aqueous ZnSO_4_ (Table S9).

The increase in wheat uptake of input-derived Zn from poultry manure was likely due to plant uptake mechanisms occurring after 8 weeks of plant growth. Our characterization of soil sampled from the pot experiment did not show any increase in potentially available Zn and Cd species with organic amendment application (Figure 3A-D). Higher-molecular-weight DOM-bound Zn increased with amendment addition (Figure 3G), and the negative correlation with DGT-extractable Zn suggests this is unlikely to be directly available for plant uptake (see Tolu et al.(20) and Figure S12). Thus, release of Zn from amendments occured over the full growth period, likely from input-derived Zn sorbed to the soil solid phases (as indicated by WHAM, Figure S13).

This release could be due to a combination of increased microbial activity and root exudation with OM degradation.(47) Co-nutritional affects, in which increased N availability promotes increased Zn uptake, may also increase with amendments enrichment in rapidly degradable OM (e.g., Künzli et al.(23)). This is due to release of available N from OM mineralization, which likely influenced the poultry manure treatment (as 50% of applied C was respired and poultry manure treatment increased soil nitrate compared to the control; Figures S6, S8-C). Organic compounds released by plants and microbes (e.g., organic acids, siderophores) could also compete with soil surface sites to form soluble metal-organic complexes,(48,49) releasing input-derived Zn species into the soil solution from previously unavailable pools.(47,50–52) In particular, phytosiderophores released by wheat are an important strategy for Zn acquisition and likely play an essential role in controlling increased Zn release with organic amendment enrichment in rapidly degradable OM.(48,53) This is because rapidly degradable OM may increase microbial activity, leading to higher competition between plants and microbes for micronutrients. This increased competition may trigger wheat plants to increase production of phytosiderophores. Thus, it is essential to further consider how biotic processes (e.g., microbial activity, root exudates) are responsible for promoting wheat crops’ access to input-derived Zn from organic amendments enriched in rapidly degradable OM.

In contrast to Zn, uptake of input-derived Cd did not increase between harvests 1 and 2 and showed low uptake with organic amendment application (Figure 1D). This could be further evidence of the role of phytosiderophores in wheat uptake of organic amendment-derived Zn versus Cd, as phytosiderophores are known to promote uptake of micronutrients such as Fe and Zn and not Cd.(54,55) Furthermore, water-extractable Cd in the original organic amendments was mainly bound to SEC fraction F2 (higher-molecular-weight DOM-Cd; Figure S11), which is unlikely to be directly available for plant uptake (Figure S12). Thus, the speciation of Cd in organic amendments could be another factor limiting its uptake by wheat. To verify the result that enrichment in rapidly degradable OM promotes wheat uptake of input-derived Zn but not Cd, further study is needed exploring a wider range of organic amendments and varied soil application rates compared to what was applied in this study. Moreover, a slight trend of increased input-derived Cd uptake with enrichment in rapidly degradable OM was observed (Figure 1D). Thus, it is necessary to further investigate whether application of organic amendments enriched in rapidly degradable OM leads to increased uptake of input-derived Cd in the wheat plants.

### IV.2 Little influence of amendments on wheat uptake of soil-derived Zn and Cd

Application of organic amendments increased wheat Zn and Cd uptake via direct addition, and not from the soil (Figure 1). This outcome was contrary to our hypotheses, as we expected the amendment most enriched in rapidly degradable OM (i.e., poultry manure) would solubilize soil-derived Zn and Cd as a result of OM degradation processes. We hypothesized this because previous studies have indicated the application of rapidly degradable OM increased wheat uptake of Zn and Cd from the soil solid phases.(23,51) However, these previous studies only tested green manures, which are even more enriched in rapidly degradable OM than poultry manures.(16) In addition, these previous studies performed pot experiments with soils that have a pH greater 7 and that contained inorganic carbon. The addition of green manure to alkaline soil rich in inorganic carbon decreased the pH, which could be an important mechanism leading to Zn solubilization from the soil solid phases.(51) This is in contrast to our study, as soil pH was not affected by organic amendment application (Figure S8-A). The ammonium treatment showed decreased pH compared to the control, likely due to nitrification of the NH_4_^+^ to NO_3_^−^ resulting in a net proton increase.(56) This pH drop with the ammonium treatment demonstrated that decreased soil pH led to increased wheat uptake of soil-derived Zn and Cd (Figure 1E,F), as previously reported with decreased pH.(57,58) Thus, it is likely because of the moderately acidic pH of the soil used in this study that application of amendments did not increase wheat uptake of soil-derived Zn and Cd through pH-related desorption processes.(59–61)

### IV.3 Organic amendments did not affect Zn and Cd concentrations in wheat grains

Overall, we did not find that organic amendment enrichment in rapidly degradable OM increased Zn and Cd in wheat grains (Figure 2). Our results suggest it is more effective to increase Zn concentrations with mineral Zn and N fertilizers than with organic amendments on the moderately acidic Cambisol used in this study (Figure 2A). This aligns with previous research that has shown varied effects of organic amendment application, with some studies showing increased Zn concentrations with organic amendment application,(51,52,62) and others showing no effect on grain Zn.(23,62)

It is noteworthy that increased total uptake of Zn with organic amendment application did not translate to increased grain Zn (Figures 1A, 2A). Storage and internal translocation of Zn may have been affected by amendment-derived N and P. N compounds bind Zn during internal translocation,(63,64) and both N and P-compounds play a role in storage of Zn in tissue. (64–67) Thus, changes to N and P could potentially affect the internal plant mechanisms that determine the distribution of Zn translocated to grains versus stored in shoot biomass. This is supported by previous studies showing changes in plant nutrient uptake alter plant expression of genes controlling micronutrient homeostasis.(68,69) Furthermore, the N-containing compound nicotianamine is known to play an important role in Zn translocation to grains, and is also a precursor used for synthesis of the phytosiderophore 2′-deoxymugineic acid.(70) If wheat grown with organic amendment application increased production of phytosiderophores to increase Zn uptake (see discussion point IV.1), this may have been at the expense of nicotianamine that could be used for internal translocation of Zn to grains. These results highlight that it is crucial to investigate how internal translocation processes of wheat cultivars are affected by organic amendment application.

## V. Conclusions

Stable isotope tracing showed wheat uptake of input-derived Zn increased with higher amendment OM degradability.(16) Input-derived Cd also followed this trend, though only very low uptake was observed. Soil analyses did not show greater available Zn and Cd with amendment application, regardless of enrichment in rapidly degradable OM. Thus, the increased Zn and Cd uptake with amendment enrichment in rapidly degradable OM was likely due to release mechanisms from the soil solid phases. This was supported by increased Zn and Cd sorbed to metal oxides and soil organic matter compared to the control, as determined by WHAM modelling. Our results highlight the importance of amendment chemical composition in controlling the flux of Zn and Cd into crops versus accumulating in soil. However, application of amendments did not lead to increased grain concentrations of Zn and Cd, despite increasing total plant uptake. This highlights the necessity of further studying the effect of organic amendments on the mechanisms of Zn translocation to grains. Finally, DGT-extractable Zn and Cd correlate negatively with species bound to higher-molecular-weight DOM (i.e., SEC fraction F2). This indicates the higher-molecular-weight DOM-bound species of Zn and Cd identified by SEC are likely not available for plant uptake.

## VI. Acknowledgements

We wish to thank Monika Macsai for all of her help with pot experiment planning, plant watering, pesticide spraying, and keeping the wheat alive and healthy. Many thanks to both Monika Macsai and Federica Tamburini for their help with isotope labeling of soil, and to Eva Ruedin for her help in preliminary tests of our experimental workflow. Thank you to Agroscope colleagues David Widmer, Jochen Mayer, Lucie Gunst, Hans-Ulrich Zbinden, and as well all colleagues at Agrovet-Strickhof for their help with organic amendment sampling. Thanks are due to Federica Tamburini for total CNS measurement in the soil and plant biomass. We are grateful to Laurie Mauclaire-Schoenholzer for double-distilling all concentrated HNO_3_ and HCl used in our work. Many thanks Natalie Gsteiger and the rest of the AuA team at Eawag for their help in measuring major anions in soil water extracts. We are grateful to Lisa Konrad, Fernando Brito, Florent Leyvraz, and Robin Krischer for support with analyzing samples and washing roots. We also thank Geremia Pellegri for his help with maintaining the wheat crops and monitoring their growth, in addition to E. Marie Muehe and Aleksandra Pienkowska for their helpful advice prior to the soil analysis. Thank you to the ETH research commission for providing funding for this work (Project No. 1-004915-000) and a big thanks to the groups of plant nutrition and inorganic environmental geochemistry at ETH for hosting and supporting this project.

## VII. Supplementary information

File includes detailed descriptions of methods, quality control, and analytical results complementary to the main text.

