## SupplementaryInformation for "The effect of organic amendment composition on zinc and cadmium availability and uptake in wheat crops"

**Affiliations:**

**Present addresses:**

**Method S1. Methods used for soil characterization by the company Sol-Conseils (Gland, Switzerland)**

Prior to all Sol-Conseil analyses, soil was dried for 48 h at 40 ºC and sieved to 2 mm. All soil analyses were conducted according to the methods of Agroscope Switzerland. The soil pH was measured by adding 50 mL ultrapure water to 20 g soil (1:2.5 soil mass-extractant volume ratio).[1] Cation exchange capacity was measured by adding 80 mL of a 0.1 M HCl and 0.025 M $H_2SO_4$ solution to 20 g soil (1:4 soil mass-extratant volume).[2] The difference between the initial extraction solution pH and the pH of the extractant (supernatant) was used to determine exchangeable $H^+$ concentrations, while exchangeable $K^+$, $Na^+$, $Ca^{2+}$ and $Mg^{2+}$ were determined via inductively coupled plasma atomic emission spectroscopy (ICP-AES). Soil texture was determined via granulometry after destruction of soil organic matter with $H_2O_2$.[3]

**Method S2. Acid digestion for total elemental analysis**

DTPA extract

Acid digestion of diethylenetriaminepentaacetic acid (DTPA) soil extracts was performed by adding 2 mL $HNO_3$ (65%, double-distilled from EMSURE grade) to 2 mL of DTPA extract. Digestion runs were identical to acid extraction of plant material (see below).

Soil

Soil (100 mg) and solid certified reference material (CRM) were digested in triplicates following a modified version of the protocol used in Imseng et al.[4] On a hotplate (120 °C), 4 mL $HNO_3$ (15 M) and 2 mL HF (23 M) were added to the sample and were digested for at least 48 h. Digested samples were dried, 4 mL 7 M HCl was added, and after at least 12 h (120 °C) the digested samples were evaporated to dryness. Four mL of $HNO_3$ (15 M) was added to the samples, and after at least 12 h (120 °C) they were evaporated to dryness. Then, 5 mL of 0.3 M $HNO_3$ was added, and the samples were refluxed overnight. Samples were then evaporated, and 4 mL concentrated HCl was added. Samples were allowed to sit for 12 h, evaporated to dryness, and 4 mL of concentrated $HNO_3$ was added. After 12 hours, samples were evaporated to dryness. Finally, 5 mL of 0.3 M $HNO_3$ was added, and samples were refluxed overnight.

Plant biomass

Wheat plant material (200 mg) and plant-derived CRM (WEPAL 939, Lucerne/*Medicago sativum*) were acid-digested with 2 mL of $HNO_3$ (65%, double-distilled from EMSURE grade) and 2 mL of ultrapure water (resistance > 18 MΩ), following a modified method of Yan et al.[5] The program of the single reaction chamber microwave (turboWave, MWS microwave systems) was as follows: temperature ramp from 25 to 220 ˚C over 23 min (125 bar of pressure applied) followed by 8 min at 220 ˚C (125 bar of pressure applied). The 15 mL borosilicate glass tubes used for the digestion were cleaned by soaking in a warm bath (60 ˚C) containing 12 M $HNO_3$ for at least 24 hours, followed by thorough rinsing with ultrapure water. Each digestion run included at least one procedural blank and one CRM. After digestion, the digests were diluted with ultrapure water to a total volume of 10 mL, then stored at 4°C until elemental quantification.

**Method S3. Quantification of total elements in soil and plant extracts by ICP-MS/MS**

All elements, i.e., Na, Mg, K, Ca, P, Mn, Fe, Ni, Cu, Zn, Cd, and Ba were quantified using an Agilent 8900 ICP-MS/MS. This device was equipped with a concentric nebulizer (Micromist),, a Scott double-pass spray chamber cooled to 2 °C, a high-throughput injection system (ISIS) with a PTFE sample loop, and platinum sampler and skimmer cones. All ICP-MS/MS parameters were optimized using a tuning solution containing 10 µg $L^{-1}$ of lithium (Li), yttrium (Y), cobalt (Co), cerium (Ce), and tellurium (Te) (prepared with standards from the company J.T. Baker). The collision gases and isotopes measured are provided in Table S1. With He collision gas, measurements were performed in single-quadrupole mode with 4.5 mL $min^{-1}$ He. With $O_2$ collision gas, measurements were performed in MS/MS mode with 5 mL $min^{-1}$ $H_2$ and 30% $O_2$. Acquisition parameters were: 0.05-0.3 ms integration time (depending on element) and 3 replicates.

For all elements, quantification was performed by external calibration using standards from Bernd Kraft (Multi-element-standard 21 elements, multi-anion-standard 5 elements) and Sigma-Aldrich (VI). The calibration standards were prepared in 1% $HNO_3$ (i.e., the same matrix as the analyzed samples). An internal standard containing scandium (Sc; 450 µg $L^{-1}$) and lutetium (Lu; 45 µg $L^{-1}$) was injected with the sample into the ICP-MS/MS to monitor instrument sensitivity throughout the measurement.

**Table S1. Quality control for elemental quantification.** Measured and certified elemental concentrations (average ± 1 standard deviation, n = 12 extraction replicates), recoveries (%) and absolute errors (%) for the digested and analyzed solid CRM WEPAL 939 (lucerne), validating the procedure of quantification of elements in wheat biomass by digestion and ICP-MS/MS analysis. For samples measured with $O_2$ collision gas, the target element isotope mass is provided and "→" indicates the actual measured isotope (e.g., for P and S this is 16 mass units higher than the element due to targeting of ions with one oxygen atom added).

| Element | Collision gas | Isotope | Units | WEPAL 939 (lucerne) [element]$_{measured}$ av ± 1 sd | WEPAL 939 (lucerne) [element]$_{certified}$ av ± 1 sd | Recovery (%) | Abs. error (%) |
|---|---|---|---|---|---|---|---|
| Na | He | 23 | g (kg)$^{-1}$ | 1.19 ± 0.05 | 1.11 ± 0.09 | 107 ± 5 | 7 ± 4 |
| Mg | He | 24 | g (kg)$^{-1}$ | 2.5 ± 0.1 | 2.4 ± 0.1 | 103 ± 4 | 4 ± 3 |
| K | He | 39 | g (kg)$^{-1}$ | 18.1 ± 0.7 | 17.2 ± 1.0 | 106 ± 4 | 6 ± 3 |
| Ca | He | 43 | g (kg)$^{-1}$ | 14.4 ± 1.0 | 13.5 ± 0.8 | 105 ± 7 | 6 ± 6 |
| P | O2 | 31 → 47 | g (kg)$^{-1}$ | 2.5 ± 0.1 | 2.5 ± 0.1 | 101 ± 4 | 3 ± 2 |
| S | O2 | 34 → 50 | g (kg)$^{-1}$ | 2.6 ± 0.2 | 2.6 ± 0.2 | 102 ± 6 | 5 ± 3 |
| Mn | He | 55 | mg (kg)$^{-1}$ | 75 ± 4 | 70.4 ± 4.4 | 105 ± 5 | 6 ± 4 |
| Fe | He | 56 | mg (kg)$^{-1}$ | 245 ± 16 | 230 ± 24 | 107 ± 7 | 7 ± 6 |
| Cu | He | 63 | mg (kg)$^{-1}$ | 5.1 ± 0.3 | 5.1 ± 0.6 | 100 ± 5 | 4 ± 2 |
| Zn | He | 66 | mg (kg)$^{-1}$ | 72 ± 4 | 65 ± 4 | 109 ± 6 | 10 ± 6 |
| Cd | O2 | 111 → 111 | mg (kg)$^{-1}$ | 0.47 ± 0.03 | 0.46 ± 0.03 | 103 ± 7 | 6 ± 4 |
| Ba | He | 137 | mg (kg)$^{-1}$ | 9 ± 1 | 8 ± 1 | 115 ± 14 | 18 ± 9 |

**Table S2. Details on the organic amendments applied in this study, along with information on the source, storage, and sampling period.**

| Organic input type | Sample name | Source | Storage/composting time | Sampling period |
|---|---|---|---|---|
| Monogastric farmyard manure | Poultry manure | Agrovet-Strickhof (Zurich, Switzerland) | Fresh from stalls (adult chickens) | October 2020 |
| Ruminant farmyard manure | Cattle manure | Agroscope-Reckenholz (Zurich, Switzerland) | <1 month in manure pile | November 2020 |
| Industrial compost | Compost | Bio massehof (Winterthur, Switzerland) | Three months of composting | November 2020 |

### Note S1. Previously published characterization of organic matter and Zn and Cd speciation in a wide range of organic amendments.

The current experiment was designed to build upon our previous organic amendment characterization study. In our previous study, a wide range of organic amendments were sampled, including green manures (n=7), lignified crop residues and lignin (n=4), monogastric farmyard manures including poultry and pig manures (n=4), ruminant farmyard manures including cattle manure (n=6), cattle slurry (n=3), and industrial compost (n=4) were collected, for a total of n=28 samples.

We analyzed organic matter molecular composition using pyrolysis gas chromatography coupled to mass spectrometry (Py-GC/MS). Using hierarchical cluster analysis, we showed that organic amendment OM molecular composition generally followed the categories of amendments sampled ("Figure 1" from Bachelder et al.).[6] Of the amendments sampled, the organic amendments most likely to be relatively enriched in rapidly degradable organic matter were green manures and lignified crop residues and litter, followed by monogastric farmyard manure. Compared to these three amendment categories, ruminant farmyard manures were relatively depleted in rapidly degradable OM. The samples most depleted in rapidly degradable OM were industrial compost samples.

We also characterized speciation of water-soluble Zn and Cd in the organic amendments ("Table 2" from Bachelder et al.). Speciation did not follow organic amendment type as closely as organic matter molecular composition. We identified an overall trend of increased Zn bound to lower-molecular-weight dissolved OM in organic amendments enriched in rapidly degradable OM, compared to amendments depleted in rapidly degradable OM. In contrast, Cd was mainly bound to higher-molecular-weight OM in all organic amendments, regardless of enrichment vs. depletion in rapidly degradable OM. In ruminant farmyard manures and slurries (e.g., cattle manure), Zn and Cd were primarily in the free $Zn^{2+}$ and $Cd^{2+}$ form, whereas the major species was DOM-bound in all other organic amendments. For more detailed discussion of our characterization, please refer to our previous publication in *Environmental Science & Technology.*[6]

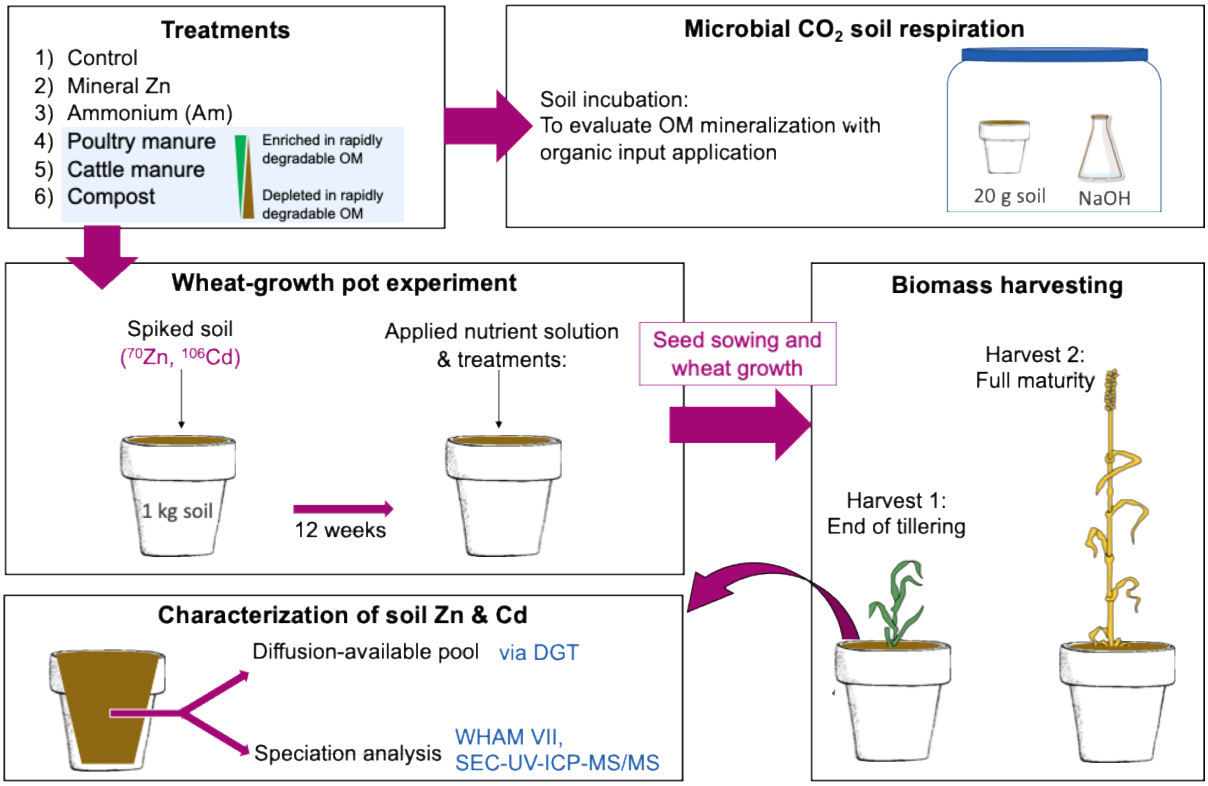

**Figure S1. Schematic overview of wheat-growth pot experiment setup, biomass harvesting, pot soil characterization, and microbial $CO_2$ respiration soil incubation.** Biomass harvesting took place 8 weeks (harvest 1, end of tillering) and 19 weeks (harvest 2, full maturity) after wheat seed sowing. Pot soil characterization of diffusion-available species was performed using diffusive gradients in thin films (DGT). For speciation analysis, Zn and Cd complexes with dissolved organic matter (DOM) were measured in water extracts, using size exclusion chromatography coupled to ultraviolet detection and triple-quadrupole mass spectrometry, i.e. SEC-UV-ICP-MS/MS[7]. Multisurface modelling (WHAM VII) was also used to calculate the speciation of 0.43 M $HNO_3$-extractable Zn and Cd species.

**Table S3. Basal nutrient solution composition for the pot experiment and application rate of select nutrients with organic amendment application.**

| | | Control | Mineral Zn, $ZnSO_4$ | Ammonium, $(NH_4)_2SO_4$ | Poultry manure | Cattle manure | Compost |
|---|---|---|---|---|---|---|---|
| Nutrient solution | N-$NH_4$, mg (kg soil)$^{-1}$ | 10 | 10 | | 10 | 10 | 10 |
| | N-$NO_3$, mg (kg soil)$^{-1}$ | 90 | 90 | 0 | 74.6 | 85.2 | 88.3 |
| | P, mg (kg soil)$^{-1}$ | 35 | 35 | 35 | 35 | 35 | 35 |
| | S, mg (kg soil)$^{-1}$ | 54 | 54 | 168[a] | 54 | 54 | 54 |
| | K, mg (kg soil)$^{-1}$ | 50 | 50 | 50 | 50 | 50 | 50 |
| | Mg, mg (kg soil)$^{-1}$ | 25 | 25 | 25 | 25 | 25 | 25 |
| | Ca, mg (kg soil)$^{-1}$ | 143 | 143 | 143 | 115 | 131 | 136 |
| Treatments | C, g (kg soil)$^{-1}$ | | | | 2 | 2 | 2 |
| | N, mg (kg DW soil)$^{-1}$ | | | | 242 | 162 | 164 |
| | N-$NH_4$, mg (kg DW soil)$^{-1}$ | | | 100 | 13.6 | 1.8 | 0.9 |
| | N-$NO_3$, mg (kg DW soil)$^{-1}$ | | | | 0 | 3 | 0.8 |
| | Zn, mg (kg DW soil)$^{-1}$ | | 1.5 | | 1.1 | 0.7 | 1.8 |
| | Cd, $\mu$g (kg DW soil)$^{-1}$ | | | | 2.2 | 1.2 | 4.4 |
| | Total mass added, g (kg DW soil)$^{-1}$ | | | | 6.2 | 6.4 | 13 |

[a]Added extra due to S content in $(NH_4)_2SO_4$

**Method S4. Greenhouse conditions, plant protection, and wheat biomass harvesting in pot experiment.**

Seeds of wheat (*Triticum aestivum,* cv. "Bobwhite") were soaked in a 30 % $H_2O_2$ solution for 15 minutes. Seeds were rinsed with ultrapure water and germinated in wet 2 mm sand that was triple washed with ultrapure water. Seedlings were sown three days after soil application of treatments and nutrient solution. During plant growth, pot soil was maintained at 30-70% WHC. Pots were randomized once per week.

The wheat was grown in a greenhouse for 4 weeks, after which it was moved to a growth chamber due to an infestation of powdery mildew. In the greenhouse, the wheat was grown with a photoperiod of 14 h (23 ºC, 65 % relative humidity) and a 10 h darkness period (20 ºC, 60 % relative humidity). In the growth chamber, the wheat was grown with a 13 h photoperiod (24º C, 55 % relative humidity, illuminance of 25 klx) and an 11 h darkness period (18 ºC, 50 % relative humidity). Due to the powdery mildew infestation, pesticides were sprayed 4 weeks after sowing. The pesticides were composed of ultrapure water containing 0.5 mL $(L)^{-1}$ Input (active agents: spiroxamine, Prothioconazole) and 0.3 mL $(L)^{-1}$ Etalfix Pro (active agent: polyether modified trisiloxane). In the sprayed solution, the total solution concentrations were 114 µg $(L)^{-1}$ Zn and 1.5 µg $(L)^{-1}$ Cd). At each application, a total of 300 mL of pesticide solution sprayed on all plants (i.e, in total 34 µg Zn and 0.5 µg Cd were sprayed on all pots, the equivalent of 0.6 µg Zn $(kg\ soil)^{-1}$ and 0.01 µg Cd $(kg\ soil)^{-1}$).

At harvest, above-ground biomass was harvested by cutting plants 2 cm above the soil surface with ceramic scissors, followed by rinsing for 20 seconds with ultrapure water to remove dust and soil particles. Roots were gently separated from soil and washed for a total of 10 minutes at 4 ºC in three successive baths of 6 mM NaCl.[8] Both roots and shoots were frozen (-20 ºC) and freeze-dried for four days, after which their dry weights were recorded. At harvest 2, the same procedures were applied as for harvest 1, but additionally grains were separated from the rest of the dried aboveground biomass.

### Table S4. Application rate of organic amendments in incubation experiment

| | Control | Mineral Zn | Ammonium* | Poultry manure | Cattle manure | Compost |
|---|---|---|---|---|---|---|
| Total mass added, g (kg soil)$^{-1}$ | | | | 4.4 | 4.6 | 9.3 |
| Total C, g (kg soil)$^{-1}$ | | | | 1.5 | 1.5 | 1.5 |
| Total N, mg (kg soil)$^{-1}$ | | | 50 | 171 | 116 | 117 |
| $N\text{-}NH_4$, mg (kg soil)$^{-1}$ | | | 50 | 9.7 | 1.3 | 0.7 |
| $N\text{-}NO_3$, mg (kg soil)$^{-1}$ | | | | < LOD | 2.2 | 0.5 |
| Total Zn, mg (kg soil)$^{-1}$ | | 1 | | 0.7 | 0.5 | 1.3 |
| Total Cd, $\mu$g (kg soil)$^{-1}$ | | | | 1.6 | 0.8 | 3.1 |

*Added extra due to S content in $(NH_4)_2SO_4$

### Note S2. Comparison of Zn and Cd quantification in non-purified and purified samples.

Zn and Cd concentrations were measured using the **conventional method** for quantifying metals in samples i.e., non-purified samples measured as if Zn and Cd in the samples were at natural abundance (Method S3).[9,10] Zn and Cd concentrations were also measured in purified samples by performing **isotope external calibration** i.e., using the summed counts per second of mass-bias-corrected all isotope data to create a calibration curve (Method S5). A comparison of these two analyses showed that for plant shoots from harvest 1, the difference between the methods ranged from -4 to 1 % for Zn quantification and -4 to -11 % for Cd quantification. This can be compared to the error measured when analyzing WEPAL 939 CRM, where the range for (absolute) error of Zn quantification was 10 ± 6 % and the range for (absolute) error of Cd quantification was 6 ± 4 % (Table S1). These results indicated that the **conventional calibration** was acceptable for quantification of Zn and Cd in isotopically enriched samples.

**Method S5. Purification and measurement of isotope ratios, including operating conditions of ICP-MS and correction of mass biases and isobaric interferences.**

Sample purification pre-treatment

Aliquots of plant extracts were transferred to acid-cleaned PTFE beakers and mixed with 2 mL of 8 M $HNO_3$ (double-distilled from Supelco Emsure grade). Samples were evaporated to dryness on a hot plate (60 ºC). Dried samples were redissolved in 1 M HCl (technical grade, triple-distilled) and were transferred to shrink-fit PTFE mini-columns[11] filled with acid-cleaned anion exchange resin (AG 1-X8, 100–200 mesh, chloride form, Bio-Rad laboratories) pre-conditioned with 2 x 1 M HCl. Matrix elements were eluted with 2 x 1.5 mL 1 M HCl. Elution of Zn and Cd was performed with 3 x 1.5 mL 0.3 M $HNO_3$. Samples were evaporated to dryness and redissolved in 0.3 M $HNO_3$ for $^{70}Zn/^{66}Zn$ and $^{106}Cd/^{111}Cd$ measurement via ICP-MS (Method S3). With each purification batch, 15-32 samples were processed in addition to at least one blank and one natrual-abunsance reference sample for quality control. Sample blanks were evaluated to ensure they contributed <5% compared to concentrations in samples. Repeat isotope ratio analysis of a quality control sample showed relative standard deviation values below 5.5% for Zn and 2.0% for Cd.

Isotope ratio analysis

To measure isotope ratios ($^{70}Zn/^{66}Zn$, $^{106}Cd/^{111}Cd$) in all plant samples, purified samples were analyzed using an Agilent 8900 ICP-MS/MS. This device included a concentric nebulizer, a Scott double-pass spray chamber cooled to 2 °C, a high-throughput injection system (ISIS) with a PTFE sample loop, and platinum sampler and skimmer cones. To tune the ICP-MS/MS, a solution was used containing 10 µg $(L)^{-1}$ of Li, Y, Co, Ce, and Te (prepared from standards from the company J.T. Baker). Select samples were analyzed with an Agilent 7500 ICP-MS equipped with a helium octopole reaction cell (5 mL $min^{-1}$ He), a concentric nebulizer, a Scott double-pass spray chamber cooled to 2°C, and nickel sampler and skimmer cones. All stable isotopes of Zn ($^{64}Zn$, $^{66}Zn$, $^{67}Zn$, $^{68}Zn$, $^{70}Zn$) and Cd ($^{106}Cd$, $^{108}Cd$, $^{110}Cd$, $^{111}Cd$, $^{112}Cd$, $^{113}Cd$, $^{114}Cd$, $^{116}Cd$) were measured. Corrections for isobaric interferences were calculated for $^{64}Zn$ ($^{64}Ni$), $^{70}Zn$ ($^{70}Ge$), $^{106}Cd$ ($^{106}Pd$), $^{108}Cd$ ($^{108}Pd$), $^{112}Cd$ ($^{112}Sn$), $^{113}Cd$ ($^{113}In$), $^{114}Cd$ ($^{114}Sn$), and $^{116}Cd$ ($^{116}Sn$). This was done by measuring non-interfering isotopes (i.e., $^{60}Ni$, $^{72}Ge$, $^{105}Pd$, $^{115}In$, and $^{115}Sn$) and using known natural-abundance isotope ratios to calculate the amount of signal originating from each interfering isotope. Mass bias corrections were performed using the sample-bracketing method. Briefly, a standard solution (40 µg Zn $(L)^{-1}$, 2 µg Cd $(L)^{-1}$) was analyzed once every four samples. The mass bias correction factor was calculated via (S1):

$$\left(\frac{^{y}Zn}{^{66}Zn}\right) corrected = \left(\frac{^{y}Zn}{^{66}Zn}\right)_{measured} e^{-kM} \qquad \text{(S1)}$$

Equation S1 was also used for Cd, with the non-spike isotope (in the denominator) being $^{111}Cd$. In (S1), *k* is the mass bias and *M* is the mass difference (calculated as *y-66* for Zn, *y-111* for Cd). The averaged *k* values between the standards measured before and after the samples were used to correct each sample in the bracket. Quantification of Zn and Cd was performed by measuring an external standard calibration (1-100 µg $(L)^{-1}$ Zn, 0.2-20 µg $(L)^{-1}$ Cd). Standards of known concentrations of Zn and Cd were purified and analyzed in parallel to samples, with recovery of Zn found to be 105±5 % and recovery of Cd found to be 98±2%. The abundances of $^{66}Zn$, $^{70}Zn$, $^{111}Cd$, and $^{106}Cd$ were calculated in the control treatment plants. The equations derived by McBeath et al were used to calculate the fractions of Zn and derived from isotopically enriched soil versus from non-enriched amendments (Eq. 3, 4) and the use efficiency (Eq. 5, 6).[9,12] These equations require calculation of the isotope abundance (*A*) of the native ($^{66}Zn$, $^{111}Cd$) and spike ($^{70}Zn$, $^{106}Cd$) isotopes, as well as the total Zn in the plant digest ($Zn_{sink}$, $Cd_{sink}$) and the total Zn added with the inputs ($Zn_{added}$, $Cd_{added}$).

$$Fraction\ Zn_{df_{fertilizer}} = \frac{A_{66,soil}\ x\ (\frac{^{70}Zn}{^{66}Zn})sink - A_{70,soil}}{(A_{70,fertilizer} - A_{70,soil}) - \left(\frac{^{70}Zn}{^{66}Zn}\right)_{sink} x(A_{66,fertilizer} - A_{66,soil})}$$

$$Fraction\ Cd_{df_{fertilizer}} = \frac{A_{111,soil}\ x\ (\frac{^{106}Cd}{^{111}Cd})sink - A_{106,soil}}{(A_{106,fertilizer} - A_{106,soil}) - \left(\frac{^{106}Cd}{^{111}Cd}\right)_{sink} x(A_{111,fertilizer} - A_{111,soil})}$$

$$Fertilizer\ use\ efficiency\ (\%) = 100\ x\ \frac{Fraction\ of\ Zn_{df_{fertilizer}} x\ Zn_{sink}}{Zn_{added}}$$

$$Fertilizer\ use\ efficiency\ (\%) = 100\ x\ \frac{Fraction\ of\ Cd_{df_{fertilizer}} x\ Cd_{sink}}{Cd_{added}}$$

Quantification of Zn and Cd in isotopically enriched samples

In select samples, a test was performed in which concentrations were calculated using the counts per seconds (i.e., raw data from the ICP-MS measurement) summed accross all Zn and Cd isotopes measured compared to the method of total elemental quantification described in Method S3, after which Method S3 was used for total elemental quanitification of Zn and Cd. See Note S2 for more details.

Tissue concentrations of Zn and Cd derived from inputs vs. soil were derived by multiplying the values of $Fraction_{df\ fertilizer}$ with the total concentration of Zn and Cd in the plant tissue. The quantity of Zn and Cd taken up by plants was calculated by multiplying plant biomass weight with tissue concentrations (Table S8).

**Table S5.** Measured and certified elemental concentrations (average ± 1 standard deviation, n= 3 measurements replicates), recoveries (%) and absolute errors (%) for liquid certified reference material NIST 1643f (freshwater), validating the procedure of quantification of elements in liquid samples via ICP-MS/MS analysis.

| Element | Collision gas | Isotope | Units | NIST 1643f | | Recovery (%) | Abs. error (%) |
|---|---|---|---|---|---|---|---|
| | | | | $[element]_{measured}$ av ± 1 sd | $[element]_{certified}$ av ± 1 sd | | |
| Na | He | 23 | $\mu g\ (L)^{-1}$ | 18590 ± 744 | 18830 ± 250 | 99 ± 4 | 3 ± 2 |
| Mg | He | 24 | $\mu g\ (L)^{-1}$ | 7275 ± 242 | 7454 ± 60 | 98 ± 3 | 4 ± 1 |
| Al | He | 27 | $\mu g\ (L)^{-1}$ | 131 ± 13 | 133.8 ± 1.2 | 98 ± 10 | 8 ± 3 |
| K | He | 39 | $\mu g\ (L)^{-1}$ | 1892 ± 66 | 1932.6 ± 9.4 | 98 ± 3 | 3 ± 1 |
| Ca | He | 43 | $\mu g\ (L)^{-1}$ | 31206 ± 948 | 29430 ± 330 | 117 ± 1 | 17 ± 1 |
| Mn | He | 55 | $\mu g\ (L)^{-1}$ | 36 ± 2 | 37.14 ± 0.6 | 97 ± 5 | 5 ± 2 |
| Fe | He | 56 | $\mu g\ (L)^{-1}$ | 93 ± 4 | 93.44 ± 0.78 | 99 ± 5 | 3 ± 2 |
| Cu | He | 63 | $\mu g\ (L)^{-1}$ | 20 ± 1 | 21.66 ± 0.71 | 92 ± 5 | 8 ± 5 |
| Zn | He | 66 | $\mu g\ (L)^{-1}$ | 72 ± 4 | 74.4 ± 1.7 | 97 ± 5 | 4 ± 1 |
| Cd | O2 | 111 → 111 | $\mu g\ (L)^{-1}$ | 5.64 ± 0.04 | 5.89 ± 0.13 | 96 ± 1 | 4 ± 1 |
| Ba | He | 137 | $\mu g\ (L)^{-1}$ | 488 ± 18 | 518.2 ± 7.3 | 94 ± 3 | 6 ± 3 |

**Note S3. Discussion of limit of detection for Zn quantification in water extracts of pot soil.**

The concentration of Zn in pot soil water extracts was extremely low, ranging from 1.9 to 18 ug $(L)^{-1}$ (Table S6). All samples were above the limit of detection (calculated as 3*standard deviation of the blank), but ten samples (of 28 in total) were below the limit of quantification (calculated as 10*standard deviation of the blank). This could introduce artefacts into our speciation measurements with SEC-UV-ICP-MS/MS.[7] This is especially true because we cannot blank-correct our data (since we measure Zn bound to organic matter, and there is no organic matter in the blank). This also shows that in this study, we are at the lower limit of water-extractable Zn concentrations in soil that can be measured with our SEC method.

**Table S6. Concentration of Zn in water extracts of soil sampled from pot experiment.** Highlighted in the table are results for water extractions of experimental (pot) replicates (rep) of soil sampled from no Zn input ("control" in main text), ammonium input, and soil in which organic amendments (poultry manure, cattle manure, and compost) were applied. The ten samples (of a total 28) that had concentrations below the limit of quantification of our method are presented, though they were above the limit od detection. This data shows that some of our samples were close to the limit of quantification for soil water-extractable Zn.

| | Zn $\mu$g $(L)^{-1}$ |
|---|---|
| Limit of detection | 1.10 |
| Limit of quantification | 2.95 |
| No Zn (rep 3) | 1.96 |
| No Zn (rep 4) | 2.25 |
| Ammonium (rep 3) | 2.77 |
| Ammonium (rep 4) | 2.75 |
| Poultry manure (rep 1) | 1.95 |
| Poultry manure (rep 2) | 1.68 |
| Poultry manure (rep 3) | 2.21 |
| Poultry manure (rep 4) | 1.90 |
| Cattel manure (rep 1) | 2.45 |
| Compost (rep 2) | 2.75 |

**Method S6. SEC-UV-ICP-MS/MS analysis of pot soil water-extracts**

To perform size exclusion chromatography coupled to ultraviolet detection and inductively coupled plasma tandem mass spectrometry (SEC-UV-ICP-MS/MS), an Agilent 1260 Infinity II high performance liquid chromatography (HPLC) system was coupled to an Agilent diode array detector (UV, 254 nm) and an Agilent 8900 ICP-MS/MS. ICP-MS/MS operating conditions were as described in Method S3. The SEC columns included one Shodex OH-pak SB-803 column and one SB-802.5 HQ column (separation <100 kDa and ~40 nm). The mobile phase used for SEC separation was ammonium nitrate (5 mmol $L^{-1}$, pH 8, flow rate of 1 mL $min^{-1}$, injection volume of 100 µL).

Each water extract was measured by SEC-UV-ICP-MS/MS twice using different gases in the collision/reaction cell (C/RC) to obtain speciation data on Zn and Cd together with various elements. Firstly, 5 mL $min^{-1}$ He gas was used in the C/RC to detect Zn (*m/z* 66; 0.3 ms integration time) and Fe (m/z 56->56; 0.1 ms). Secondly, 25% $O_2$ and 1 mL $min^{-1}$ $H_2$ was used in the C/RC to detect P, S, Zn, and Cd. An internal standard containing Sc (450 µg $L^{-1}$) and Lu (45 µg $L^{-1}$) was injected into the ICP-MS/MS with the sample to monitor instrument sensitivity throughout the measurement period. The chromatograms collected for Zn speciation with He and O2 collision gasses were the same. Thus, data analysis was performed using data collected with $O_2$ due to higher sensitivity. Semi-quantitative speciation data for Zn, Cd, P, S, Fe, and UV ($Abs_{254nm}$) was obtained by deconvoluting the intensity chromatograms of these elements by using Origin 2021 (Fit peak Pro), as outlined by Laborda et al.[13]

The recovery of the Zn and Cd in the SEC peaks compared to the total Zn and Cd in the water extracts was determined via direct injection and peak integration using the software MassHunter. Recovery was calculated as the total SEC peak area divided by the total peak area of the direct injection peak (i.e., injection of the sample into the ICP-MS/MS without first passing it through the SEC column).

**Note S4. Water-extracted fractions in SEC-UV-ICP-MS/MS data**

We applied our previously optimized method to analyze water extracts of soil and organic amendments. This application of size exclusion chromatography coupled to ultraviolet detection and inductively coupled plasma tandem mass spectrometry (SEC-UV-ICP-MS/MS) has been discussed extensively in our previous publications.[6,7] For details on the optimization of operating conditions for this technique, we refer readers to the supplementary information sections of our previous work. Generally, this method allows separation and characterization of distinct size and chemical fractions of dissolved organic matter and nanoparticles <40 nm in diameter. For compound separation by SEC, size separation occurs with larger molecules eluting at shorter retention times and smaller molecules eluting at higher retention times. Proof of this separation based on size is provided (Figure S2-A). Compounds are also separated by chemical properties due to secondary interactions between analyte species and the column stationary phase, with more negatively charged molecules eluting earlier and less negatively charged molecules eluting later. Thus, fractions with earlier retention times comprise higher-molecular-weight, more negatively charged molecules and fractions with shorter retention times comprise lower-molecular-weight, less negatively charged molecules. To demonstrate that our separation based on size and chemical properties separates distinct fractions of OM, we analyzed isolated and purified standards from the international humic substances society (IHSS). As expected, we found that the higher-molecular-weight, more negatively charged humic acid standard eluted earlier than the lower-molecular-weight, less negatively charged fulvic acid standard (Figure S2-B).

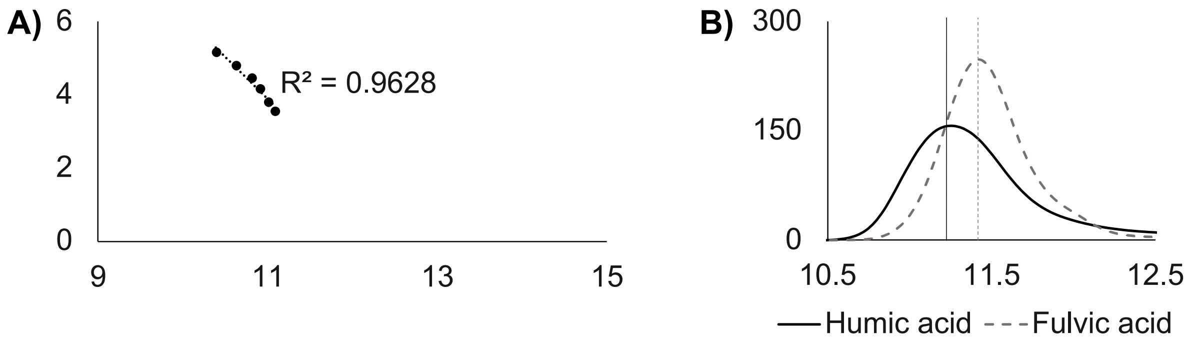

**Figure S2. Analysis of A) molecular weight and B) humic substance reference standards to validate separation by size & chemical composition using SEC-UV-ICP-MS/MS.** Panel A) provides log correlation of molecular weight standards (polypropylene sulfonate) with their retention time. Panel B) provides Elliott soil reference standard from international humic substances society, with a dotted line showing fulvic acid and a solid line showing humic acid references. Vertical lines indicate the retention times of each peak apex.

**Note S4 (cont.). Water-extracted fractions in SEC-UV-ICP-MS/MS data**

The chromatograms of UV and all elements measured were highly similar between treatments and replications. Figure S3 provides a sample chromatogram including UV, P, S, Fe, Zn, and Cd. Based on these chromatograms, three fractions of distinct DOM size and chemical composition were defined. F1 was associated with peaks of Fe and P, with a small amount of UV signal as well (Figure S5). This fraction eluted before reference OM materials (F1 peak apex < 10.8 mins, humic substance elution >11.2 mins). Fraction F1 also had very low UV signal. Thus, it is likely that F1 represents small (oxy)hydroxide nanoparticles as previously observed for soil.[7] F2 was associated with large UV peaks, indicating high amounts of aromatic OM in this fraction. The observed sharp S peak in this fraction (elution at ~12.2 mins) was likely due to $SO_4^{2-}$, which elutes in F2 due to strong negative repulsion between this anion and the stationary phase. Fraction F2 also contained small peaks of P, either from organic P compounds or $PO_4^{3-}$ sorbed to DOM. We designated F2 as higher-molecular-weight OM because it was highly aromatic and contained peaks with similar retention times as reference OM materials from the IHSS (F2 peak apexes 10.8-12.5 mins humic substance elution 11.2-11.5 mins). F3 was characterized by UV and P peaks that with retention times of 12.5-15 mins. As fraction F3 eluted after all IHSS reference compounds, this fraction was considered to comprise lower-molecular-weight DOM.

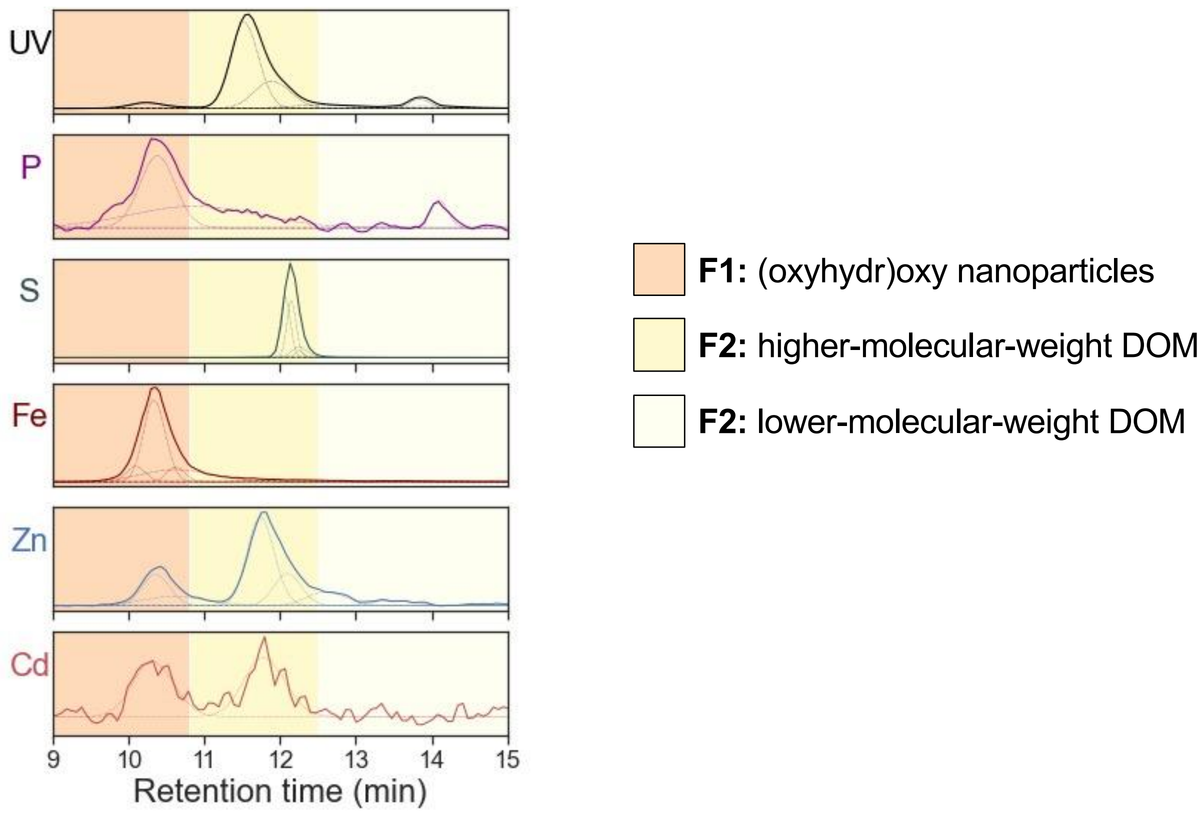

**Figure S3. Sample chromatogram from SEC-UV-ICP-MS/MS analysis of water-extracted pot soil (control treatment).** Figure shows the ultraviolet (UV), phosphorus (P), sulfur (S), iron (Fe), zinc (Zn), and cadmium (Cd) chromatograms for one experimental (pot) replicate. For all elements, the intensity chromatograms are shown (in counts $s^{-1}$) in solid lines, while dotted lines show peak deconvolutions. Shaded panels indicate peak apex time ranges of three size and chemical fractions F1, F2, and F3.

**Method S7. Chemical equilibrium modelling inputs and sensitivity tests.**
The model input parameters included pH ($H_2O$), dissolved organic carbon, dissolved major anions (input as $NO_3^-$, $SO_4^{2-}$, $F^-$ quantified in soil water extracts), carbonate ions, and reactive dissolved organic matter (DOM). Concentrations of carbonate ions were determined from the partial pressure of $CO_2$ in the atmosphere, which was set to 0.004 atm following Mossa et al.[14] Total DOM was calculated by multiplying DOC by a factor of 2 as assumed previously in WHAM modelling studies.[15,16] Of total DOM, 65% was considered "reactive" and able to bind metals.[17] For analysis of water-extractable species, total concentrations of water-extractable elements (Na, Mg, Al, Ca, Mn, Fe, Ni, Cu, Zn, Cd, Ba, Pb)) were included as inputs. Water-extractable $PO_4^{3-}$ was calculated from total dissolved P. For solid-phase speciation, model inputs included metal oxides (Fe, Mn, and Al), reactive solid-phase organic matter (SOM, i.e. non-dissolved organic matter), and total concentrations of 0.43 M $HNO_3$-extractable elements (Na, Mg, Al, K, Ca, Mn, Fe, Ni, Cu, Zn, and Cd). Metal oxides were calculated as the Fe, Mn, and Al extracted by dithionite-citrate-bicarbonate method.[18] Reactive SOM was calculated as 30% of total SOM following the results of Duffner et al.'s investigation into multisurface modelling of low-Zn soil, with SOM estimated as total soil carbon multiplied by a factor of two as assumed by Duffner et al.[19] The species output by the model included free $Zn^{2+}$ and $Cd^{2+}$, DOM-bound Zn and Cd ("bound to colloidal fulvic acid"), and Zn and Cd sorbed to SOM, metal oxides (Fe, Mn, and Al), and clay minerals. Sensitivity tests were performed to evaluate the effect of altering model parameters on speciation output, showing no difference in general trends for free $Zn^{2+}$, $Cd^{2+}$, or for Zn and Cd sorption to the major solid phases (Figures S4 & S5, Table S7).

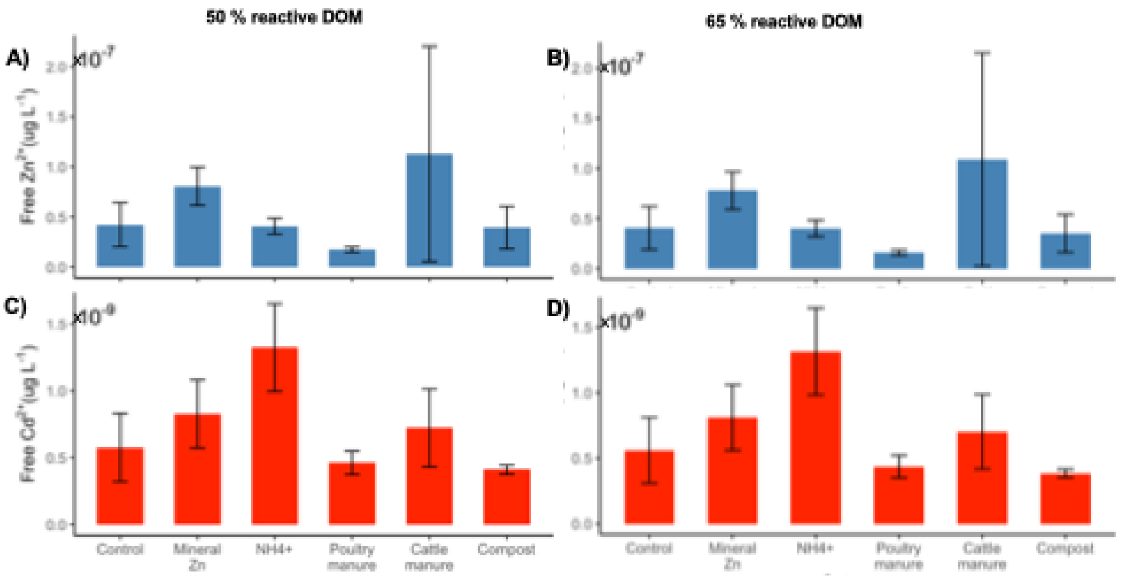

**Figure S4. Sensitivity tests of equilibrium modelling of Zn and Cd speciation in water-extractable pool.** Soil samples were collected from a wheat-growth pot experiment and extracted with ultrapure water for chemical equilibrium modelling (WHAM VII). Concentrations of water-extractable free $Zn^{2+}$ (panels A, B) and free $Cd^{2+}$ (panels C, D) with two scenarios of reactive DOM: 50% of DOM considered reactive (left panels; A, C) and 65% of DOM considered reactive (right panels; B, D). Error bars represent ±1 standard deviation from the mean as calculated from experimental (pot) replicates.

**Table S7. Summary of scenarios used for multisurface modelling (WHAM VII) sensitivity tests.** Reactive dissolved organic matter (DOM) was calculated as either 50 % or 65 % of total DOM. Reactive solid-phase organic matter (SOM) was calculated as either 15 % or 30 % of total SOM. The percentage of mineral oxides with active sorption sites that can sorb trace elements was calculated as 100 % of the total crystalline and amorphous-extractable mineral oxides (DCB-extractable minus the fractions in the 0.43 M $HNO_3$-extractable pool). In scenario 4, a mineral oxide sensitivity test (100*) was performed considering all DCB-extractable mineral oxides as containing active sorption sites (including the 0.43 M $HNO_3$-extractable pool).

| | **Scenario No.** | | | |
|---|---|---|---|---|
| **Input parameter** | **1** | **2** | **3** | **4** |
| Reactive DOM (%) | 65 | 65 | 50 | 65 |
| Reactive SOM (%) | 30 | 15 | 30 | 30 |
| Metal oxides (%) | 100 | 100 | 100 | 100* |

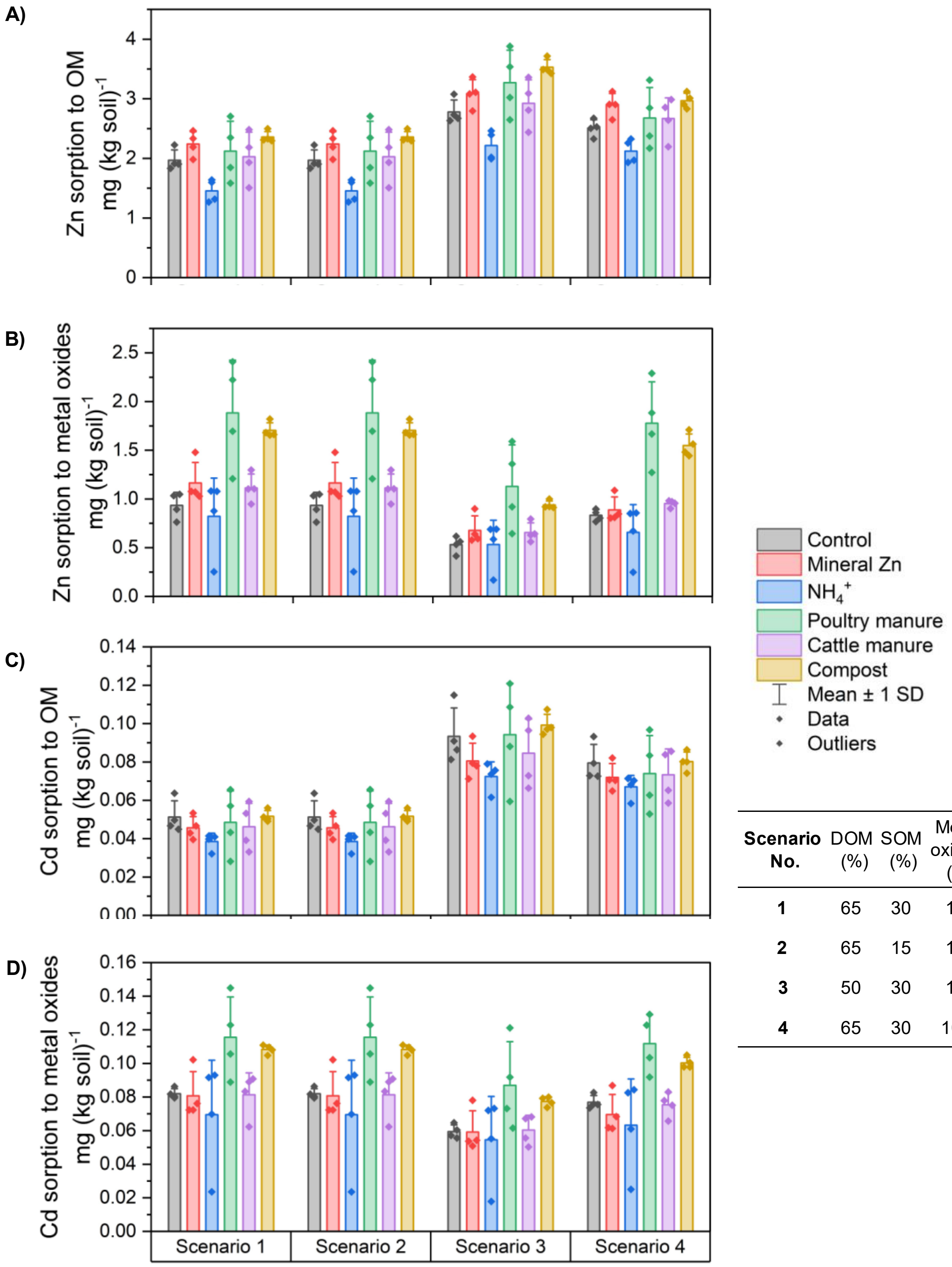

| Scenario No. | DOM (%) | SOM (%) | Metal oxides (%) |
|---|---|---|---|
| 1 | 65 | 30 | 100 |
| 2 | 65 | 15 | 100 |
| 3 | 50 | 30 | 100 |
| 4 | 65 | 30 | 100* |

**Figure S5. Sensitivity tests of equilibrium modelling of Zn and Cd speciation in 0.43 M $HNO_3$-extractable pool.** Soil samples were collected from a wheat-growth pot experiment and extracted with 0.43 M $HNO_3$ for speciation modelling of free ion and solid-phase adsorbed species. Concentrations of Zn sorption to organic matter (OM) and to metal oxides (Panels A, B) and Cd sorption (Panels C, D) were determined using the model WHAM VII. Calculations were performed for four scenarios, which are provided in Table S7 (also provided in the figure, for reference). Error bars represent ±1 standard deviation from the mean as calculated from experimental (pot) replicates.

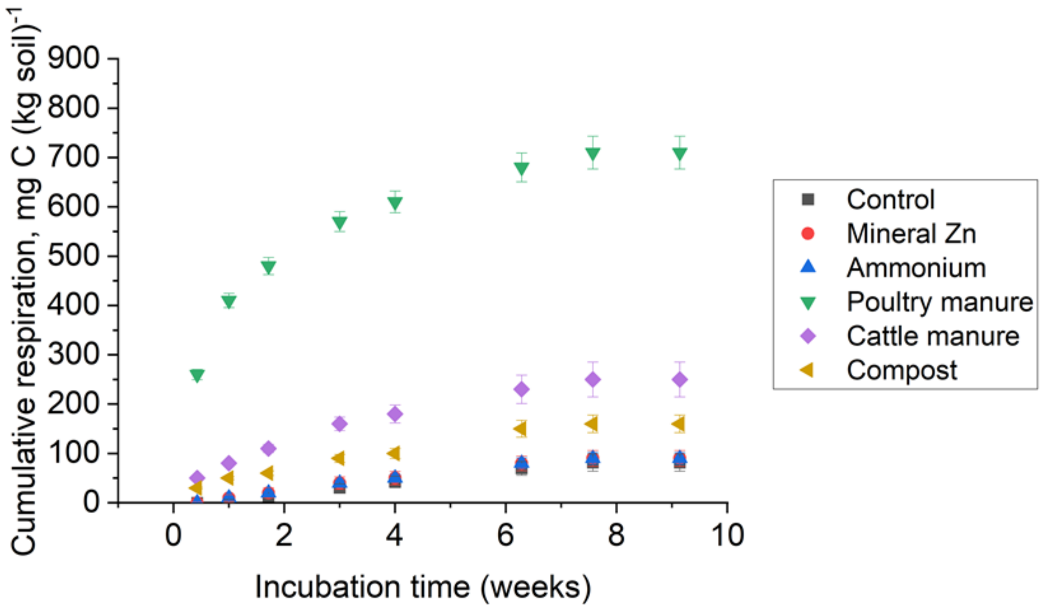

**Figure S6. Microbial $CO_2$ respiration with organic amendment application to soil.** Values indicate cumulative carbon respired over the incubation period. Error bars represent ±1 standard deviation as calculated from n=5 experimental (incubation) replicates.

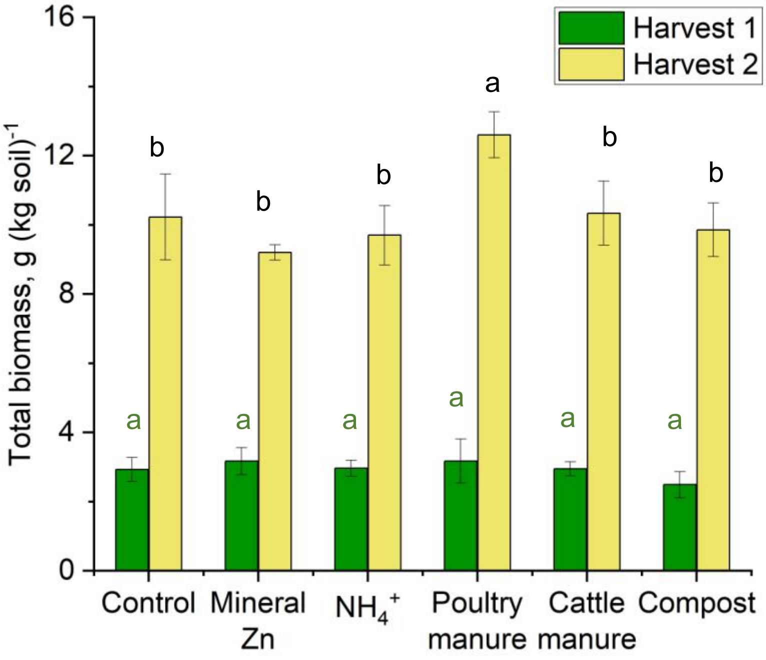

**Figure S7. Whole-plant biomass production in wheat crops harvested at harvest 1 (green) and harvest 2 (yellow).** Treatments under which wheat was grown included no Zn fertilizer (control), mineral Zn applied as aqueous $ZnSO_4$, an ammonium ($NH_4^+$) treatment applied as $(NH_4)_2SO_4$, and three organic amendments. Error bars show ± 1 standard deviation of the mean, calculated from n=4 experimental (pot) replicates. Lowercase letters indicate statistical differences between treatments at harvest 1 (eight weeks after sowing, end of tillering; green) and harvest 2 (19 weeks after sowing, full maturity; black).

**Table S8. Biomass production and total concentrations of C, N, Zn, and Cd in all plant organs.** Samples were collected at harvests 1 (H1, 8 weeks after sowing, end of tillering) and harvest 2 (H2, 19 weeks after sowing, full manturity). Error shows ± 1 standard deviation around the mean, calculated from n=4 experimental (pot) replicates. All values were calculated using the dry weight of plant biomass.

| | | Control | Mineral Zn | Ammonium | Poultry manure | Cattle manure | Compost |
|---|---|---|---|---|---|---|---|
| Biomass, g (pot)$^{-1}$ | | av ± 1sd | av ± 1sd | av ± 1sd | av ± 1sd | av ± 1sd | av ± 1sd |
| | Stem + leaves | 1.9 ± 0.2 | 2.1 ± 0.4 | 1.8 ± 0.2 | 2.2 ± 0.2 | 2.1 ± 0.1 | 1.6 ± 0.2 |
| | Root | 1.1 ± 0.2 | 1.1 ± 0.1 | 1.2 ± 0.4 | 1.0 ± 0.5 | 0.9 ± 0.1 | 0.9 ± 0.2 |
| H2 | Grain | 3.8 ± 0.6 | 3.1 ± 0.2 | 3.4 ± 0.4 | 5.2 ± 0.4 | 4.2 ± 0.6 | 4.2 ± 0.3 |
| | Stem + leaves[a] | 5.9 ± 0.7 | 5.6 ± 0.0 | 5.9 ± 0.4 | 6.6 ± 0.2 | 5.6 ± 0.4 | 5.1 ± 0.5 |
| | Root | 0.5 ± 0.1 | 0.5 ± 0.1 | 0.5 ± 0.1 | 0.8 ± 0.2 | 0.5 ± 0.1 | 0.6 ± 0.2 |
| Total C, g (kg plant)$^{-1}$ | | | | | | | |
| H1 | Stem + leaves | 410 ± 1 | 410 ± 2 | 407 ± 2 | 409 ± 1 | 408 ± 1 | 410 ± 1 |
| | Root | 390 ± 7 | 399 ± 8 | 389 ± 7 | 397 ± 8 | 388 ± 3 | 391 ± 2 |
| H2 | Grain | 406 ± 2 | 408 ± 0 | 407 ± 1 | 408 ± 1 | 408 ± 1 | 406 ± 1 |
| | Stem + leaves[a] | 416 ± 2 | 418 ± 2 | 410 ± 1 | 418 ± 2 | 414 ± 2 | 416 ± 1 |
| | Root | 439 ± 1 | 441 ± 2 | 439 ± 2 | 433 ± 1 | 437 ± 2 | 440 ± 2 |
| Total N, g (kg plant)$^{-1}$ | | | | | | | |
| H1 | Stem + leaves | 35 ± 2 | 32 ± 2 | 37 ± 1 | 38.6 ± 0.5 | 39 ± 1 | 36 ± 1 |
| | Root | 19.6 ± 0.3 | 20 ± 1 | 20 ± 1 | 20 ± 1 | 21 ± 1 | 19.6 ± 0.3 |
| H2 | Grain | 24 ± 1 | 27 ± 2 | 27 ± 2 | 26 ± 2 | 26 ± 3 | 22 ± 1 |
| | Stem + leaves[a] | 7 ± 1 | 7 ± 1 | 9 ± 1 | 8 ± 1 | 7 ± 1 | 6.2 ± 0.4 |
| | Root | 13 ± 1 | 13.6 ± 0.2 | 13 ± 1 | 14.9 ± 0.3 | 13.3 ± 0.3 | 14 ± 1 |
| Total Zn, mg (kg plant)$^{-1}$ | | | | | | | |
| H1 | Stem + leaves | 27 ± 2 | 30 ± 2 | 35 ± 4 | 29 ± 4 | 34 ± 3 | 27 ± 2 |
| | Root | 19 ± 2 | 22 ± 1 | 26 ± 2 | 22 ± 2 | 25 ± 2 | 19 ± 1 |
| H2 | Grain | 29 ± 3 | 38 ± 3 | 38 ± 4 | 31 ± 5 | 34 ± 4 | 25 ± 1 |
| | Stem + leaves[a] | 9 ± 1 | 10 ± 2 | 16 ± 2 | 8 ± 1 | 10 ± 2 | 7 ± 1 |
| | Root | 10 ± 0 | 13 ± 2 | 11 ± 1 | 7 ± 4 | 9 ± 1 | 10 ± 1 |
| Total Cd, µg (kg plant)$^{-1}$ | | | | | | | |
| H1 | Stem + leaves | 533 ± 81 | 460 ± 103 | 787 ± 105 | 516 ± 41 | 653 ± 81 | 466 ± 59 |
| | Root | 1'756 ± 193 | 1'530 ± 141 | 2'430 ± 197 | 1'915 ± 128 | 1'866 ± 221 | 1'422 ± 114 |
| H2 | Grain | 127 ± 30 | 104 ± 15 | 172 ± 26 | 150 ± 31 | 129 ± 9 | 104 ± 10 |
| | Stem + leaves[a] | 502 ± 48 | 339 ± 59 | 771 ± 170 | 473 ± 41 | 526 ± 32 | 405 ± 44 |
| | Root | 1'193 ± 196 | 929 ± 104 | 1'197 ± 26 | 999 ± 151 | 910 ± 63 | 925 ± 162 |

[a]Value includes all above-ground biomass except grains

**Table S9. Percentage of Zn and Cd derived from inputs ($Zn_{dff}$, $Cd_{dff}$) that was recovered in plant biomass at harvest 2.** Fertilizer use efficiencies are calculated as the amount of Zn (or Cd) in the whole plant divided by the Zn (or Cd) application rate, per kg of soil. Some values were below the limit of detection of our method (<LOD). Error bars show ± 1 standard deviation around the mean, calculated from n=4 experimental (pot) replicates. Lowercase letters (a-c) indicate statistical differences.

| | **Mineral Zn** | **Poultry manure** | **Cattle manure** | **Compost** |
|---|---|---|---|---|
| Zn application rate, mg (kg soil)$^{-1}$ | 1.5 | 1.1 | 0.7 | 1.8 |
| Whole-plant $Zn_{dff}$ (% recovery) | 5 ± 1 b | 8 ± 1 a | 7 ± 1 a | 2.0 ± 0.2 c |
| Cd application rate, $\mu$g (kg soil)$^{-1}$ | | 2.2 | 1.2 | 4.4 |
| Whole-plant $Cd_{dff}$ (% recovery) | | 17 ± 4 | 11 ± 14 | 0.2 ± 2.5 |

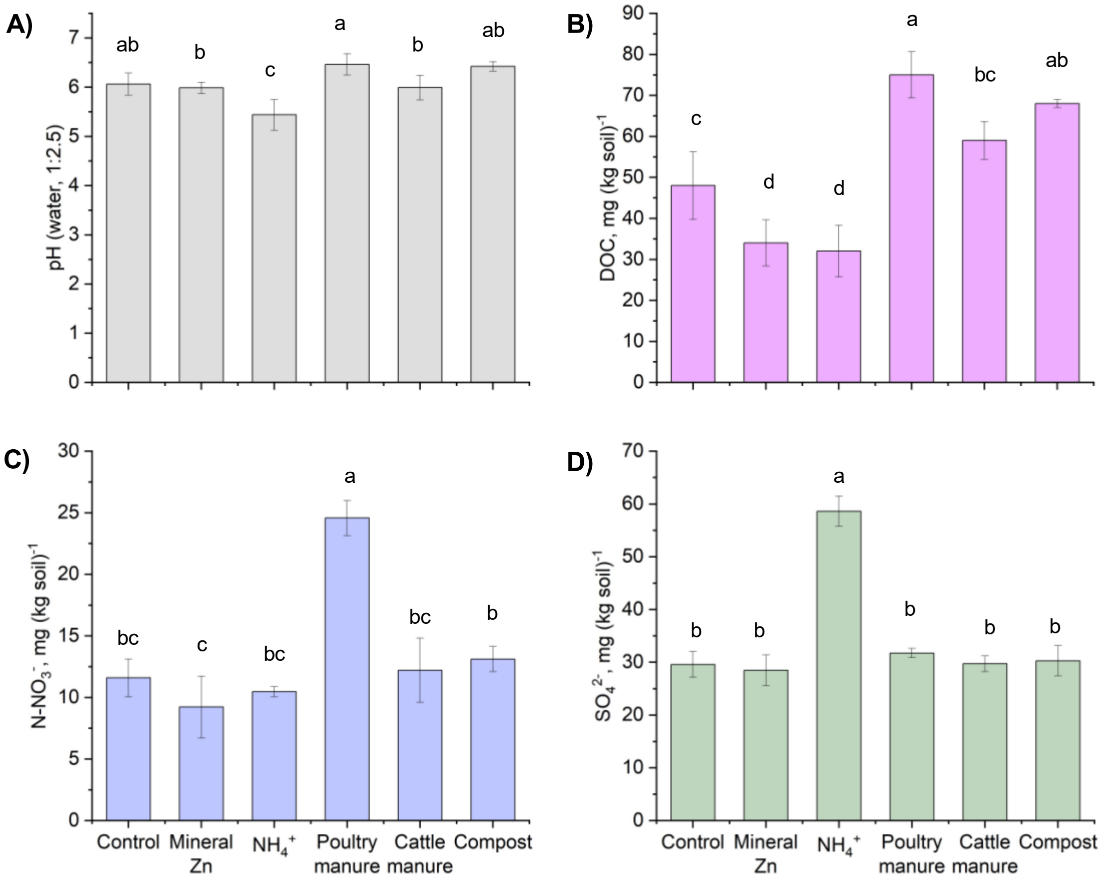

**Figure S8. Chemical composition of pot soil sampled at harvest 1 (8 weeks after wheat sowing).** Panels show A) pH and B) dissolved organic carbon (DOC) in pot soil. Treatments under which wheat was grown included no Zn fertilizer (control), mineral Zn applied as aqueous $ZnSO_4$, an ammonium ($NH_4^+$) treatment applied as $(NH_4)_2SO_4$, and three organic amendments. Error bars show ± 1 standard deviation around the mean, calculated from n=4 experimental (pot) replicates. Lowercase letters (a-d) indicate statistical differences between treatments.

**Note S5. Characterization of soil pH, DOC, $NO_3^-$, and $SO_4^{2-}$**

We evaluated how our treatments affected bulk geochemical parameters that could affect soil Zn and Cd availability. At harvest 1, the soil pH of the control, mineral Zn, and organic amendment treatments were not significantly different (Figure S8-A). However, the ammonium treatment significantly decreased soil pH to 5.4±0.3 compared to 6.0±0.2 in the control. Poultry manure and compost significantly increased the DOC to 75±6 mg C (kg soil)$^{-1}$ and 68±1 mg C (kg soil)$^{-1}$ in these treatments, respectively, compared to 48±8 mg C (kg soil)$^{-1}$ in the control (Figure S8-B). In contrast, mineral Zn and ammonium treatments decreased DOC to 34±6 mg C (kg soil)$^{-1}$ and 32±6 mg C (kg soil)$^{-1}$, respectively. Cattle manure did not lead to a significant difference in DOC compared to the control. Poultry manure application doubled $NO_3^-$ to 24±2 mg N-$NO_3^-$ (kg soil)$^{-1}$ compared to 12±2 mg N-$NO_3^-$ (kg soil)$^{-1}$ in the control (Figure S8-C). No other inputs significantly affected $NO_3^-$ compared to the control. Similar to $NO_3^-$, only the ammonium treatment increased $SO_4^{2-}$, to 59±4 mg (kg soil)$^{-1}$ compared to 29±3 mg (kg soil)$^{-1}$ in the control (Figure S8-D). This two-fold increase was likely due to the $NH_4^+$ form applied, which was $(NH_4)_2SO_4$.

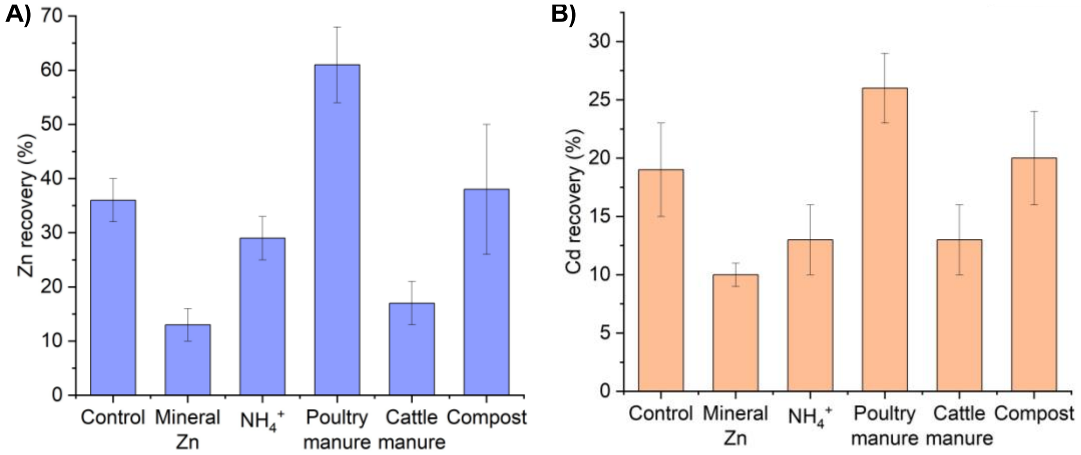

**Figure S9. Recovery of species measured by SEC-UV-ICP-MS/MS in water-extracted pot soil.** Data shows the percentage of total water-extracted A) Zn and B) Cd present in all identified SEC fractions (fractions F1-3, Figure S9). Values were calculated as the summed peak area for all SEC peaks determined with divided by the total peak area after direct sample injection (i.e. without first passing through the SEC column before ICP-MS/MS measurement). Error bars show ± 1 standard deviation around the mean, calculated from n=4 experimental (pot) replicates.

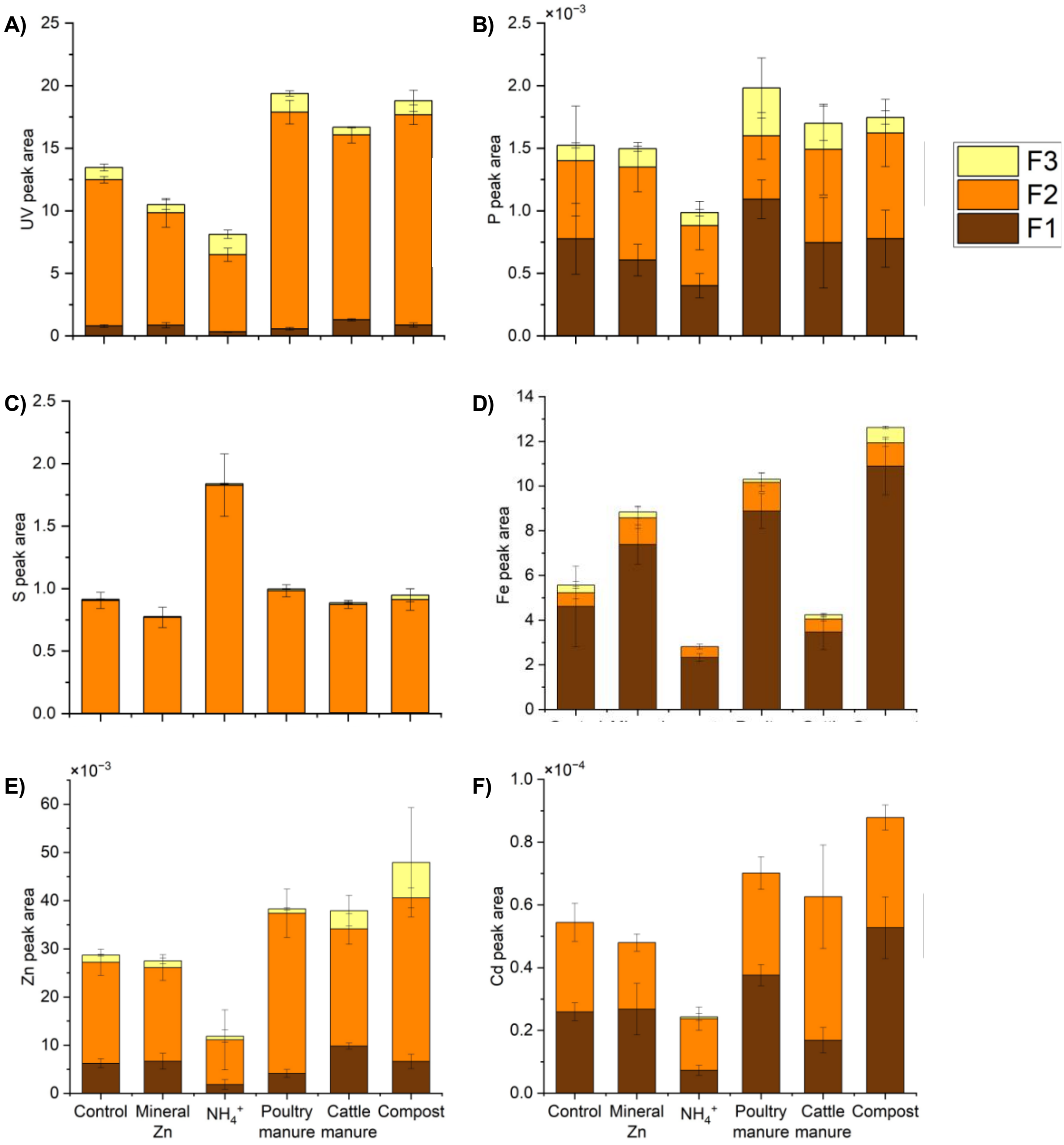

**Figure S10. Characterization of Zn and Cd associated with dissolved organic matter (DOM) in pot soil as measured by SEC-UV-ICP-MS/MS**. Bar charts show the distribution of A) ultraviolet (UV) absorption at 254 nm, B) P, C) S, D) Fe, E) Zn, and F) Cd between three size and chemical fractions: **(F1)** (organo)mineral nanoparticles, **(F2)** higher-molecular-weight dissolved organic matter (HMW DOM), and **(F3)** lower-molecular-weight dissolved organic matter (LMW DOM). Data is presented as total peak area in each fraction as determined by peak deconvolution with Origin Pro. Treatments (x-axis) include the control, mineral Zn, ammonium (Am), sunflower (SF), poultry manure (poultry), cattle manure (cattle), and compost. Error bars show ± 1 standard deviation around the mean, calculated from n=4 experimental (pot) replicates.

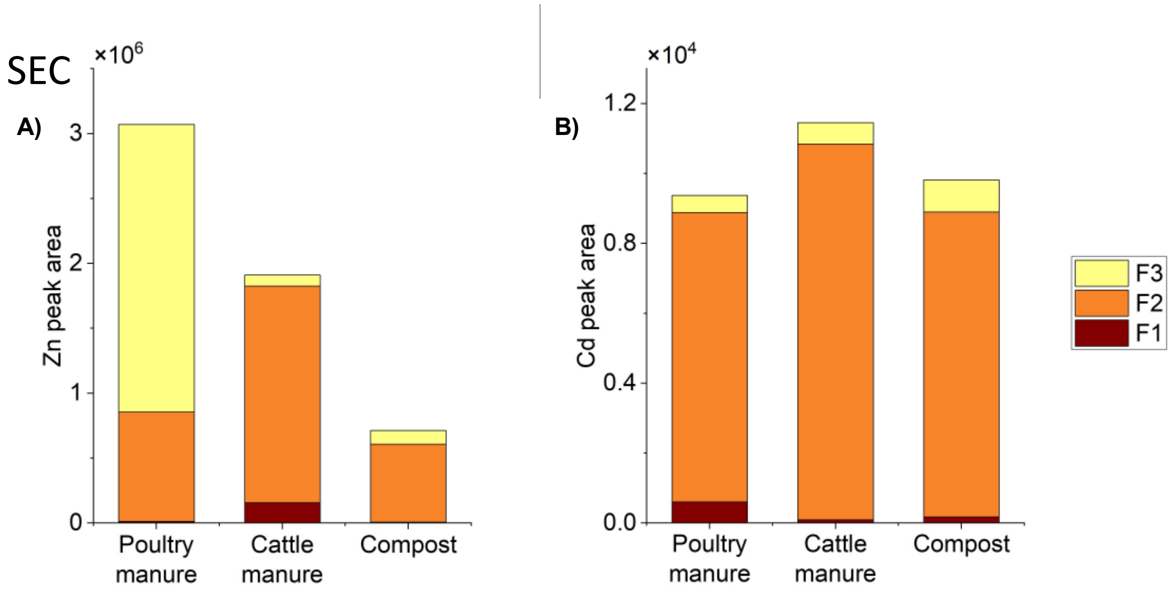

**Figure S11. Characterization of Zn and Cd speciation in original organic amendments.** Association of A) Zn and B) Cd with dissolved organic matter (DOM) in organic amendments as measured by SEC-UV-ICP-MS/MS. Bar charts show the distribution of A) Zn and B) Cd between three size and chemical fractions: **(F1)** (organo)mineral nanoparticles, **(F2)** higher-molecular-weight dissolved organic matter (HMW DOM), and **(F3)** lower-molecular-weight dissolved organic matter (LMW DOM). Data is presented as relative abundance in each fraction as determined by peak deconvolution with Origin Pro.

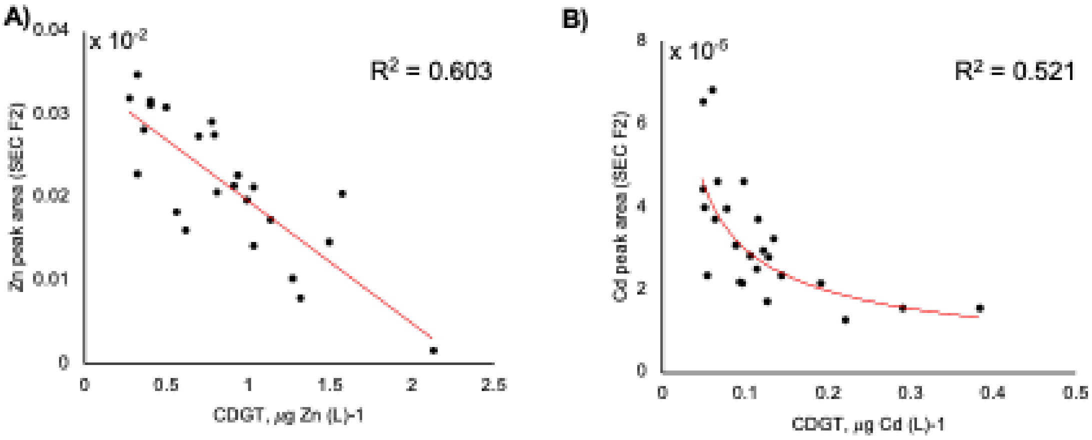

**Figure S12.** Correlations between DGT-extractable pools and SEC fraction F2 for A) Zn and B) Cd with coefficient of determination ($R^2$) values. Interpolation equations were SEC = -0.0001(DGT_Zn) + 5x$10^{-05}$ for Zn and SEC = 7x$10^{-06}$x(DGT_Cd)$^{-0.605}$ for Cd, determined using Excel.

**Table S10.** Pearson's coefficients (R values) for correlations between select soil and plant parameters. Data for size exclusion chromatography (SEC) peak areas for fractions 1, 2 and 3 (F1-3) are included for Zn and Cd. Effective DGT concentrations (C_DGT) and nitrogen nutrient indices (NNI) are also included. All data was collected at harvest 1, with the exception of parameters labeled harvest 2 (H2) and grain concentrations of Zn and Cd.

| | pH | DOC | C-DGT, Zn | C-DGT, Cd | Water-Zn | Water-Cd | CO2 resp. | SEC F1 (Zn) | SEC F2 (Zn) | SEC F3 (Zn) | SEC F1 (Cd) | SEC F2 (Cd) | Free Zn | Free Cd | Zn Uptake (H2) | Cd Uptake (H2) | Grain Zn conc. | Grain Cd conc. | NNI (H2) |
|---|---|---|---|---|---|---|---|---|---|---|---|---|---|---|---|---|---|---|---|
| **pH** | | 0.6 | 0.5 | 0.7 | 0.0 | 0.6 | 0.2 | 0.1 | 0.8 | 0.3 | 0.1 | 0.6 | 0.1 | 0.6 | 0.0 | 0.2 | 0.2 | 0.1 | 0.0 |
| **DOC** | | | 0.5 | 0.4 | 0.1 | 0.4 | 0.3 | 0.1 | 0.7 | 0.2 | 0.1 | 0.6 | 0.1 | 0.5 | 0.0 | 0.0 | 0.3 | 0.0 | 0.0 |
| **C-DGT, Zn** | | | | 0.7 | 0.1 | 0.8 | 0.2 | 0.1 | **0.7** | 0.2 | 0.1 | 0.5 | 0.1 | 0.7 | 0.1 | 0.0 | 0.3 | 0.0 | 0.0 |
| **C-DGT, Cd** | | | | | 0.0 | 0.9 | 0.2 | 0.3 | 0.8 | 0.2 | 0.3 | **0.4** | 0.0 | **0.8** | 0.1 | 0.3 | 0.2 | 0.2 | 0.0 |
| **Water-Zn** | | | | | | 0.1 | 0.0 | 0.1 | 0.0 | 0.0 | 0.1 | 0.1 | 0.4 | 0.1 | 0.1 | 0.3 | 0.0 | 0.3 | 0.2 |
| **Water-Cd** | | | | | | | 0.1 | 0.2 | 0.7 | 0.2 | 0.2 | 0.4 | 0.1 | 0.9 | 0.1 | 0.1 | 0.2 | 0.1 | 0.0 |
| **CO2_64days** | | | | | | | | 0.1 | 0.4 | 0.0 | 0.1 | 0.3 | 0.0 | 0.2 | 0.3 | 0.2 | 0.4 | 0.1 | 0.3 |
| **SEC F1 (Zn)** | | | | | | | | | 0.2 | 0.0 | 0.9 | 0.0 | 0.1 | 0.2 | 0.2 | 0.4 | 0.1 | 0.3 | 0.1 |
| **SEC F2 (Zn)** | | | | | | | | | | 0.3 | 0.2 | 0.7 | 0.0 | 0.7 | 0.1 | 0.2 | 0.3 | 0.1 | 0.0 |
| **SEC F3 (Zn)** | | | | | | | | | | | 0.0 | 0.3 | 0.1 | 0.2 | 0.0 | 0.0 | 0.0 | 0.0 | 0.0 |
| **SEC F1 (Cd)** | | | | | | | | | | | | 0.0 | 0.1 | 0.2 | 0.3 | 0.4 | 0.1 | 0.4 | 0.2 |
| **SEC F2 (Cd)** | | | | | | | | | | | | | 0.1 | 0.5 | 0.0 | 0.0 | 0.3 | 0.0 | 0.0 |
| **Free Zn** | | | | | | | | | | | | | | 0.0 | 0.0 | 0.0 | 0.0 | 0.0 | 0.1 |
| **Free Cd** | | | | | | | | | | | | | | | 0.1 | 0.2 | 0.1 | 0.2 | 0.1 |
| **Zn Uptake (H2)** | | | | | | | | | | | | | | | | 0.6 | 0.3 | 0.5 | 0.7 |
| **Cd Uptake (H2)** | | | | | | | | | | | | | | | | | 0.1 | 0.5 | 0.3 |
| **Grain Zn conc.** | | | | | | | | | | | | | | | | | | 0.2 | 0.3 |
| **Grain Cd conc.** | | | | | | | | | | | | | | | | | | | 0.4 |
| **NNI (H2)** | | | | | | | | | | | | | | | | | | | |

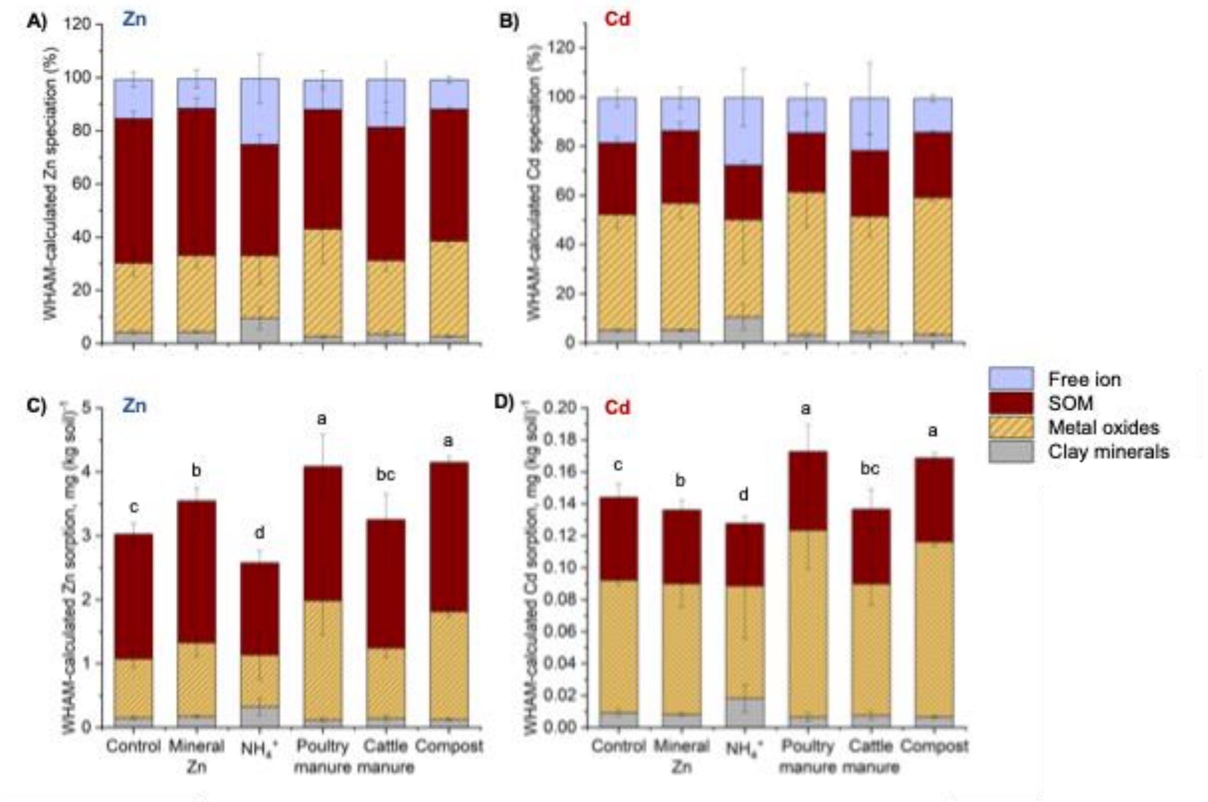

**Figure S13**. Multisurface modelling results for percentage of A) Zn and B) Cd species. Concentrations of C) Zn and D) Cd species sorbed to soil solid phases are also provided. Modelling was performed using the 0.43 M $HNO_3$-extractable Zn and Cd concentrations and includes sorption to solid-phase organic matter (SOM), metal oxides, and clay minerals. Modeling scenario 1 results are displayed, i.e. 30% total SOM considered as reactive, 65% total DOM considered as reactive, and 100% dithionite-citrate-buffer-extractable metal oxides considered as reactive (Table S7). Error bars show ±1 standard deviation of the mean, calculated from n=4 experimental (pot) replicates. Lowercase letters (a-d) indicate statistical differences between treatments for the sum of solid-phase sorbed species.

(13) Laborda, F.; Bolea, E.; Górriz, M. P.; Martín-Ruiz, M. P.; Ruiz-Beguería, S.; Castillo, J. R. A Speciation Methodology to Study the Contributions of Humic-like and Fulvic-like Acids to the Mobilization of Metals from Compost Using Size Exclusion Chromatography–Ultraviolet Absorption–Inductively Coupled Plasma Mass Spectrometry and

Deconvolution Analysis. *Analytica Chimica Acta* **2008**, *606* (1), 1–8. https://doi.org/10.1016/j.aca.2007.10.048.
(14) Mossa, A.-W.; Gashu, D.; Broadley, M. R.; Dunham, S. J.; McGrath, S. P.; Bailey, E. H.; Young, S. D. The Effect of Soil Properties on Zinc Lability and Solubility in Soils of Ethiopia – an Isotopic Dilution Study. *SOIL* **2021**, *7* (1), 255–268. https://doi.org/10.5194/soil-7-255-2021.
(15) Lofts, S.; Tipping, E.; Hamilton-Taylor, J. The Chemical Speciation of Fe(III) in Freshwaters. *Aquat Geochem* **2008**, *14* (4), 337–358. https://doi.org/10.1007/s10498-008-9040-5.
(16) Dwane, G. C.; Tipping, E. Testing a Humic Speciation Model by Titration of Copper-Amended Natural Waters. *Environment International* **1998**, *24* (5–6), 609–616. https://doi.org/10.1016/S0160-4120(98)00046-4.
(17) Bryan, S. E.; Tipping, E.; Hamilton-Taylor, J. Comparison of Measured and Modelled Copper Binding by Natural Organic Matter in Freshwaters. *Comparative Biochemistry and Physiology Part C: Toxicology & Pharmacology* **2002**, *133* (1–2), 37–49. https://doi.org/10.1016/S1532-0456(02)00083-2.
(18) Mehra, O. P.; Jackson, M. L. Iron Oxide Removal from Soils and Clays by a Dithionite-Citrate System Buffered with Sodium Bicarbonate. *Clays clay miner. (Natl. Conf. Clays Clay Miner.)* **1960**, *7*, 317–327. https://doi.org/10.1346/CCMN.1958.0070122.
(19) Duffner, A.; Weng, L.; Hoffland, E.; van der Zee, S. E. A. T. M. Multi-Surface Modeling To Predict Free Zinc Ion Concentrations in Low-Zinc Soils. *Environ. Sci. Technol.* **2014**, *48* (10), 5700–5708. https://doi.org/10.1021/es500257e.
